# L-bodies are novel RNA-protein condensates driving RNA transport in *Xenopus* oocytes

**DOI:** 10.1101/2020.05.08.084814

**Authors:** Christopher R. Neil, Samantha P. Jeschonek, Sarah E. Cabral, Liam C. O’Connell, Erin A. Powrie, Timothy A. Wood, Kimberly L. Mowry

## Abstract

RNP granules are membrane-less compartments within cells, formed by phase separation, that function as regulatory hubs for diverse biological processes. However, the mechanisms by which RNAs and proteins interact to promote RNP granule structure and function *in vivo* remain unclear. In *Xenopus laevis* oocytes, maternal mRNAs are transported as large RNPs to the vegetal hemisphere of the developing oocyte, where local translation is critical for proper embryonic patterning. Here, we demonstrate that vegetal transport RNPs represent a new class of cytoplasmic RNP granule, termed Localization-bodies (L-bodies). We show that L-bodies are multiphase RNP granules, containing a dynamic protein-containing phase surrounding a non-dynamic RNA-containing substructure. Our results support a role for RNA as a critical scaffold component within these RNP granules and suggest that *cis*-elements within localized mRNAs may drive subcellular RNA localization through control over phase behavior.

## Introduction

Spatial organization of the cell cytoplasm has emerged as a key strategy for gene regulation. For example, reorganization of otherwise ubiquitously distributed mRNA into specific cytoplasmic foci, termed ribonucleoprotein (RNP) granules, is now recognized as a widespread post-transcriptional regulatory strategy (reviewed in Banani et al. 2017; Shin and Brangwynne 2017; Gomes and Shorter 2019). Classes of RNP granules, such as stress granules, P-bodies, neuronal transport granules, and germline specifying granules, have been generally classified by their cellular context and presumed biological function. While physically distinct, these cytoplasmic RNP granules share a striking number of common features, including both extensive compositional conservation and potential overlap in underlying biological function. In addition, emerging research on the physical characteristics of RNP granules may suggest organizational similarities between classes of RNP granules (reviewed in Banani et al. 2017).

Phase separation has emerged as a fundamental physical property of RNP granules. Multivalent RNA binding proteins (RBPs) and proteins that contain intrinsically disordered regions (IDRs) are enriched in nearly all known cytoplasmic RNP granule classes (Kedersha et al. 2013; Buchan 2014). Such proteins are thought to work in combination with RNA species that contain multiple protein-binding sites (Li et al. 2012; Han et al. 2012; Kato et al. 2012; Lin et al. 2015; Elbaum-Garfinkle et al. 2015; Nott et al. 2015; Protter et al. 2018) to drive multivalent interactions leading to phase separation (reviewed in Brangwynne et al. 2015). *In vivo*, RNP granule components are dispersed in the cytoplasm but can assemble into a condensed liquid or gel-like phase to form RNP granules in response to dynamic cues (Weber and Brangwynne 2012; Toretsky and Wright 2014). While it remains an open question as to how RNAs and proteins determine the physical characteristics of biological condensates, similarities in both composition and behavior points to RNP granules as conserved centers for post-transcriptional gene regulation.

In addition to RNP granule formation, spatial control of mRNA distribution in the cytoplasm is regulated through RNA localization (reviewed in Medioni et al. 2012; Eliscovich et al. 2013; Holt and Schuman 2013). RNA localization, which functions to generate cellular polarity in a wide variety of cell types and organisms, proceeds through the binding of *cis*-acting RNA sequences, termed localization elements, by *trans*-acting protein factors (reviewed in Suter 2018). In this way, combinations of RNA *cis*-elements and RBPs are required for assembly of specific RNPs that can be localized to distinct regions of the cytoplasm. While certain components of localized RNPs have been identified, the physical nature of these assemblies, and the mechanisms driving their formation, remain largely unknown.

*Xenopus laevis* oocytes are an important model system for the study of RNA localization. Here, mRNAs encoding germ layer determinants become restricted in the vegetal hemisphere of developing oocytes, where they are subsequently transported to the vegetal cortex and act to pattern the embryo following fertilization (reviewed in Medioni et al. 2012). mRNAs localized through this pathway, including *vegT, trim36*, and, most prominently, *vg1* (Weeks and Melton 1987; Birsoy et al., 2006; Zhang et al., 1998; Cuykendall and Houston, 2009), rely on interactions with a core set of RBPs, including hnRNPAB, PTB, Staufen, and Vera to drive the formation of transport-competent RNP structures (Deshler et al. 1997; Gautreau et al. 1997; Havin et al. 1998; Cote et al. 1999; Yoon and Mowry 2004; Czaplinski et al. 2005; Lewis et al. 2004). Transport of these RNPs to the vegetal cortex is achieved through active transport along microtubules, mediated by kinesin and dynein motor proteins (Betley et al. 2004; Messitt et al. 2008; Gagnon et al. 2013). However, the molecular and physical nature of these and other RNP transport cargos remains uncharacterized.

In this work, we have identified Localization-bodies (L-bodies) as novel biomolecular condensates. Our results indicate that L-bodies are multiphase RNP granules, composed of a non-dynamic mRNA-containing substructure enmeshed in a dynamic protein-containing layer. L-bodies are unexpectedly large and contain a heterogeneous population of localized mRNAs, packaged by RBPs and surrounded by microtubules and molecular motors. We also show that incorporation of mRNAs into L-bodies is correlated with localization and appears to rely on selective entrapment, specified by sequence features of resident mRNAs. Biochemical isolation and proteomic analysis of L-bodies reveals a high degree of compositional similarity to other classes of cytoplasmic RNP granules, with the majority of identified proteins showing either direct conservation with other RNP granule types, or a shared enrichment of multivalent RNA binding domains and IDRs. For the first time, we have characterized the composition and biophysical properties of L-bodies, providing new insights into developmentally programmed transport of maternally loaded mRNAs and the roles that RNP granules play in governing post-transcriptional gene regulation. Importantly, these results suggest a novel role for mRNA in cytoplasmic granule organization.

## Results

### L-bodies are large RNP complexes that contain vegetally localizing mRNAs

To characterize the molecular features of RNA transport cargos, we first visualized localized and non-localized mRNAs in immature *Xenopus* oocytes by fluorescence *in situ* hybridization (FISH). During stages II-III of oogenesis, vegetally localized mRNAs including *vg1, vegT*, and *trim36* are enriched in the vegetal oocyte cytoplasm during transport to the vegetal cortex (Figs. 1A, A′, S1A, A′). At higher magnification, *vg1* mRNA foci are restricted to large clusters (Figure 1B), which also contain *vegT* and *trim36* mRNA (Figs. 1B′, S1B). These heterotypic vegetal bodies, comprised of multiple RNA foci, are surprisingly large, ∼5-10 μm (Figure 1B, B′). In contrast, neither a germinal granule mRNA, *nos1*, nor a non-localized mRNA, *gapdh*, are enriched in the vegetal RNA bodies. *nos1*, having been localized earlier in oogenesis, is tightly deposited at the vegetal cortex (Fig. S1C), while *gapdh* is uniformly distributed throughout the oocyte cytoplasm (Fig. 1C′). Interestingly, while not enriched, *gapdh* is not excluded from the vegetal RNA clusters (Fig. 1D, D′). Our results show that during localization, vegetally transported mRNAs are restricted to non-exclusive, heterotypic bodies hereafter termed Localization-bodies, or L-bodies.

**Figure 1.**
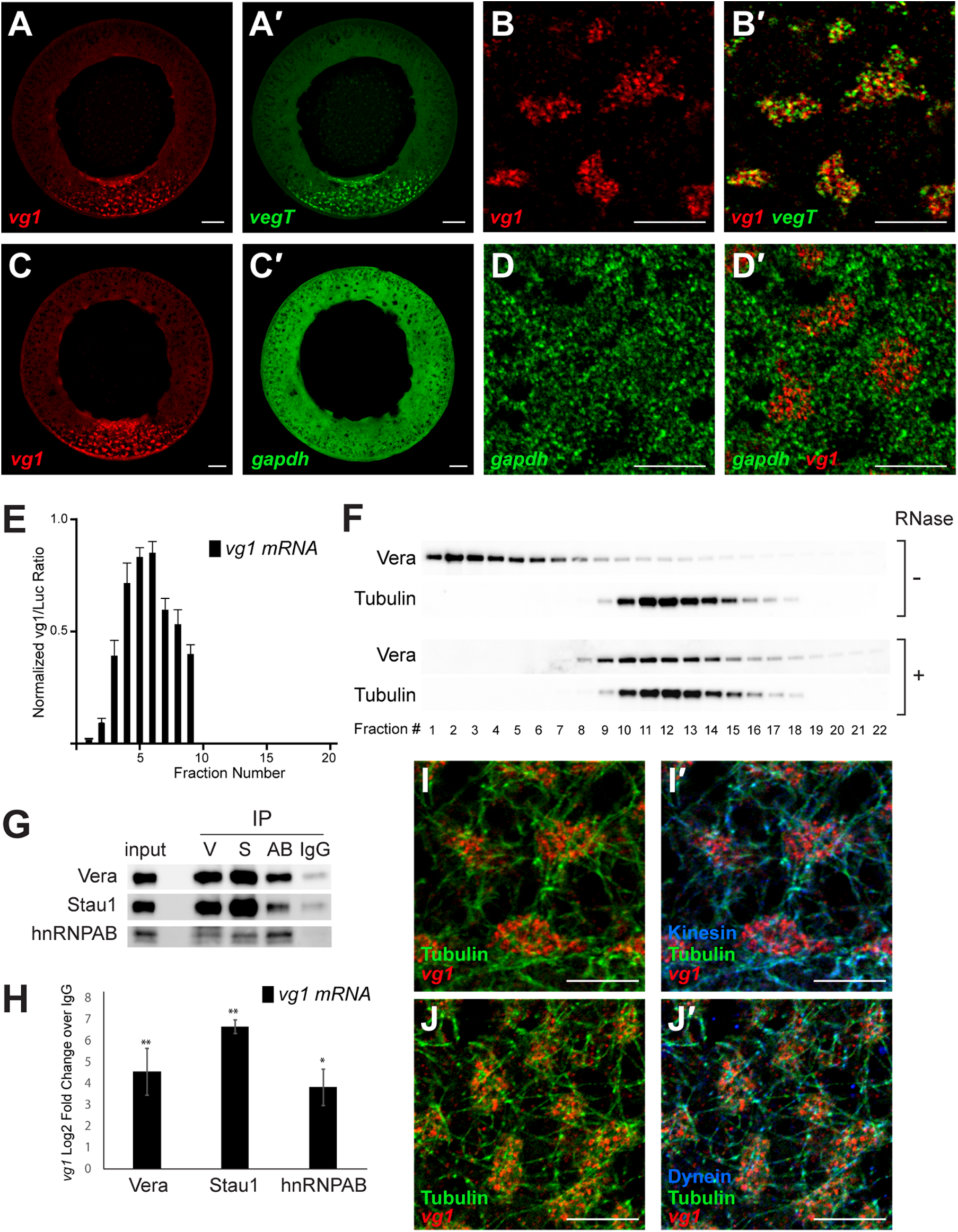
Vegetally localizing mRNA is contained in a large RNP complex. **(A)** A cryosection of a stage II oocyte probed by FISH for *vg1* mRNA (red) and *vegT* mRNA (A′, green) is shown, with the vegetal cortex at the bottom. Scale bar=50 μm. **(B)** Higher magnification view of the vegetal cytoplasm of a stage II oocyte probed by FISH for *vg1* mRNA (red), shown merged in B′ with *vegT* mRNA (green). Scale bar=10 μm. **(C)** A cryosection of a stage II oocyte probed by FISH for *vg1* mRNA (red) and *gapdh* mRNA (C′, green) is shown, with the vegetal cortex at the bottom. Scale bar=50 μm. **(D)** Higher magnification view of *gapdh* mRNA in the vegetal cytoplasm of a stage II oocyte probed by FISH for *gapdh* mRNA (green), shown merged in D′ with *vg1* mRNA (red). Scale bar=10 μm. **(E)** After size exclusion (SE) chromatography, *vg1* mRNA levels were quantitated column fractions by qRT-PCR, normalized to *luciferase* control RNA. Error bars represent standard error of the mean (*n*=11 columns). **(F)** Oocyte lysate was treated with RNase A prior to SE chromatography, followed by immunoblot analysis with anti-Vera and anti-Tubulin antibodies. Column fractions from untreated (-) lysate are shown at the top and column fractions from RNase-treated (+) lysate are shown at the bottom. **(G)** Pooled SE column fractions (2-5) were immunoprecipitated (IP) using anti-Vera (V), anti-Stau1 (S), anti-hnRNPAB (AB), and IgG. After SDS-PAGE, co-IP of Vera, Stau1, and hnRNPAB were confirmed by immunoblotting with anti-Vera (top), anti-Stau1 (middle), and anti-hnRNPAB (bottom). **(H)** Pooled SE column fractions (2-5) were IPed using anti-Vera, anti-Stau1, anti-hnRNPAB, and IgG. Following isolation of bound RNA, *vg1* RNA was detected by qRT-PCR, with normalization to a *luciferase* RNA control. Shown is log2-fold enrichment for *vg1* RNA from the Vera (left), Stau1 (middle), and hnRNPAB (right) co-IPs over IgG. *n*=5 and error bars represent standard error of mean. ** indicates p < 0.01, * indicates p < 0.05. **(I)** FISH-IF is shown for *vg1* mRNA (red) and anti-Tubulin (green), and in I′ with anti-Kinesin (blue). Scale bar=10 μm. **(J)** FISH-IF is shown for *vg1* mRNA (red) and anti-Tubulin (green), and in J′ with anti-Dynein (blue). Scale bar=10 μm. See also Figure S1.

In order to identify additional components of L-bodies, we fractionated lysates prepared from stage II-III oocytes using size exclusion (SE) chromatography. Both *vg1* mRNA (Fig. 1E), as well as the *vg1* RBP Vera (Fig. 1F), chromatograph as large complexes in the void volume. After treatment of oocyte lysate with RNase prior to SE chromatography, Vera is no longer present in the void volume and instead shifts into fractions containing smaller complexes (Fig. 1F), indicating that the large Vera-containing complexes are RNPs. By contrast, Tubulin chromatographs in fractions containing small complexes regardless of RNase treatment (Fig. 1F). Co-immunoprecipitation (co-IP) analyses using fractions pooled from the void volume after SE chromatography demonstrate that the well-characterized *vg1* RBPs Vera (Igf2bp3; Havin et al. 1998; Deshler et al. 1998), Stau1 (Staufen1; Yoon and Mowry 2004), and hnRNPAB (40LoVE; Czaplinski et al. 2005) all co-IP with one another (Fig. 1G). Importantly, these *vg1* RBPs also co-IP *vg1* mRNA from the large RNP pool (Fig. 1H). These results suggest that *vg1* mRNA, Vera, hnRNPAB, and Stau1 are packaged together in L-bodies.

We have previously shown that vegetal RNA localization relies on the microtubule motor proteins kinesin and dynein (Messitt et al. 2008; Gagnon et al. 2013). To determine whether L-bodies are associated with microtubules and motor proteins, we performed immunofluorescence in conjunction with FISH (FISH-IF) for *vg1* mRNA and Tubulin (Fig. 1I, J), kinesin 1 (Fig. 1I′) and dynein (Fig. 1J′). L-bodies are encased in microtubule baskets (Fig. 1I, J), which are decorated with both kinesin and dynein motors (Fig. 1I′, J′), suggesting that the L-body complex, rather than individual *vg1* mRNAs, are interacting with the transport machinery.

### Localized RNAs are specifically enriched in L-bodies

Localization of *vg1* mRNA can be recapitulated by a localization element (LE) that is composed of a duplication of the first 135 nucleotides of a larger element residing in the *vg1* mRNA 3′ UTR (Gautreau et al. 1997; Lewis et al. 2008). Microinjected *LE* RNA is enriched in the vegetal cytoplasm (Fig. 2A) and is both co-packaged with endogenous *vg1* mRNA in the vegetal cytoplasm (Fig. 2A′) and incorporated into L-bodies (Figure 2B, B′). By contrast, a mutant version of the *LE* RNA (*mutLE*) harboring point mutations that ablate specific RBP binding (PTB and Vera; Lewis et al. 2008) is not capable of vegetal localization (Fig. 2C) and fails to accumulate in L-bodies (Fig. 2D, D′). Incorporation of localized and non-localized RNAs into L-bodies was further examined by SE chromatography (Fig. 2E). Microinjected RNAs were tagged with unique qRT-PCR barcodes that do not affect L-body assembly (Fig. S2). As shown in Fig. 2E, both endogenous *vg1* mRNA and microinjected *LE* RNA are highly enriched in large complexes present in the void volume, while microinjected non-localized RNAs, *mutLE* and *β-globin*, are not. Taken together, these results suggest that enrichment into L-bodies is mediated by sequence specific features of target RNAs and that such enrichment is required for subsequent vegetal RNA localization.

**Figure 2.**
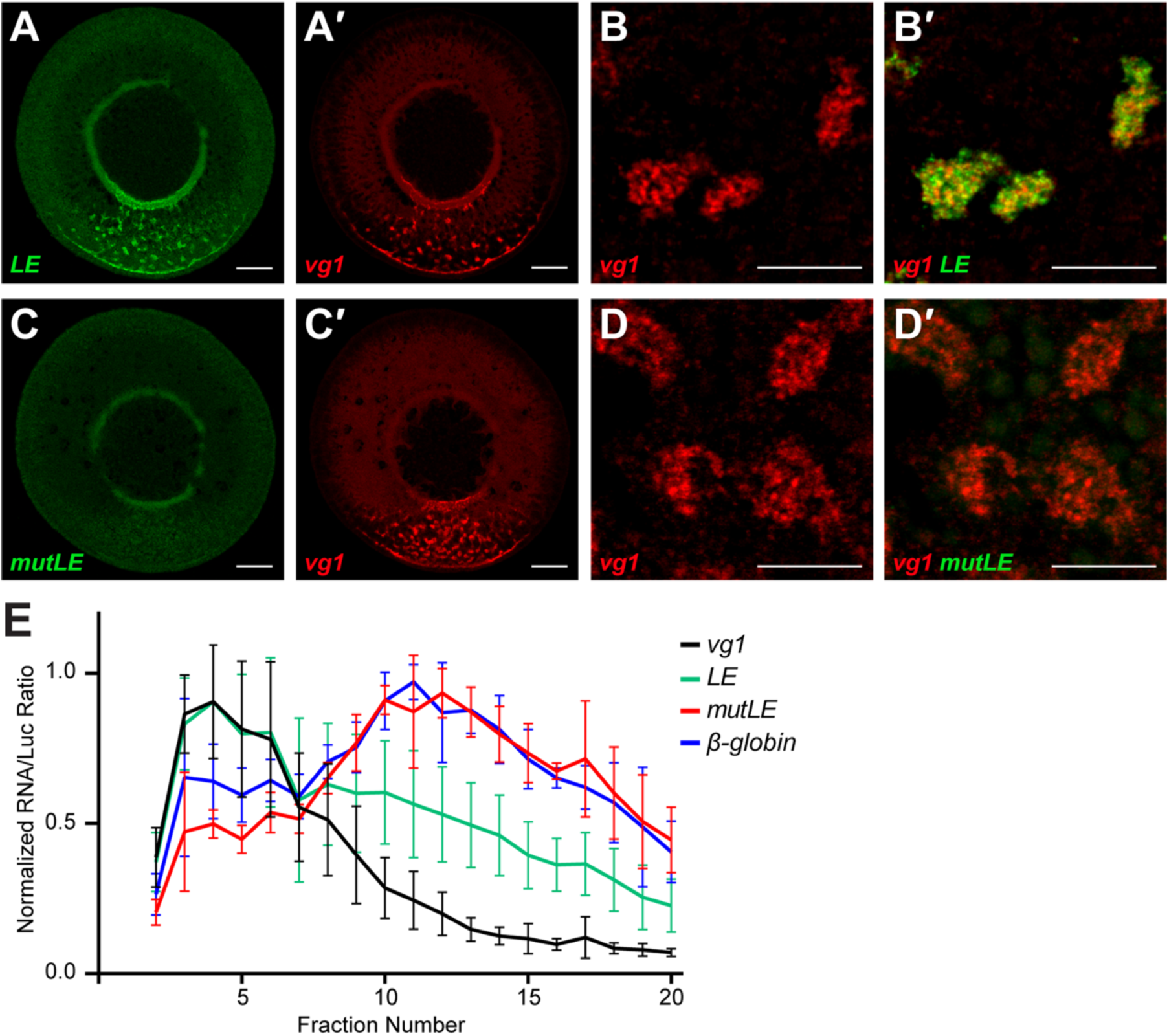
Assembly into L-bodies is correlated with vegetal localization. **(A)** A cryosection of a stage II oocyte is shown, with microinjected fluorescently-labeled *LE* RNA (green) and endogenous *vg1* mRNA (A′) detected by FISH (red) is shown. The vegetal cortex at the bottom, and the scale bar=50 μm. **(B)** High magnification view of the vegetal cytoplasm of a stage II oocyte with endogenous *vg1* mRNA (red) detected by FISH and microinjected *LE* RNA (green) shown merged in B′. Scale bar=10 μm. **(C)** A cryosection of a stage II oocyte is shown, with microinjected fluorescently-labeled mutant *LE* RNA (*mutLE*) in green and endogenous *vg1* mRNA (C′) detected by FISH in red. The vegetal cortex at the bottom and the scale bar=50 μm. **(D)** High magnification view of the vegetal cytoplasm of a stage II oocyte with endogenous *vg1* mRNA (red) detected by FISH and microinjected *mutLE* RNA (green) shown merged in D′. Scale bar=10 μm. **(E)** Stage II-III oocytes were microinjected with *LE* RNA (green), non-localizing *mutLE* RNA (red), and non-localizing *β-globin* RNA (blue). Oocyte lysates were subjected to SE chromatography and the levels of the microinjected RNAs and endogenous *vg1* mRNA (black) were quantitated in SE column fractions by qRT-PCR, normalized to *luciferase* control RNA. Error bars represent standard error of the mean (*n*=4 columns). See also Figure S2.

### Purification of L-bodies

To analyze the composition of L-bodies we took a biochemical approach (Fig. 3A). First, crosslinked oocyte lysate was enriched for large complexes by SE chromatography (Figs. 1E-F), followed by parallel immunoprecipitation (IP) using two known *vg1* RBPs, Stau1 and hnRNPAB (Fig. 3A). Stau1 and hnRNPAB, like other characterized *vg1* RBPs, are known to have multiple roles in RNA biogenesis and are not solely involved in mRNA localization (Yisraeli 2005; Snedden et al., 2013; Heraud-Farlow and Kiebler 2014). Therefore, we reasoned that requiring interactions with two different *vg1* RBPs would more accurately identify constituents specific to L-bodies. Stau1 and hnRNPAB are both highly enriched in L-bodies as assessed by FISH-IF (Fig. 3B-E) and SE chromatography (Fig. 3F). Immunoblot analysis confirmed that Stau1 and hnRNPAB co-IP with one another, but not with IgG (as in Fig. 1G), and RNA-IP followed by qRT-PCR (as in Fig. 1H) demonstrated that *vg1* mRNA was significantly enriched in the IPs compared to IgG. Liquid chromatography-tandem mass spectrometry (LC-MS/MS) was performed on each of the three IPs, requiring candidate proteins to be enriched at least 2-fold over the IgG control in both the Stau1 and hnRNPAB IPs. These results identified a set of 86 proteins that represent potential L-body components (Fig. 3G).

**Figure 3.**
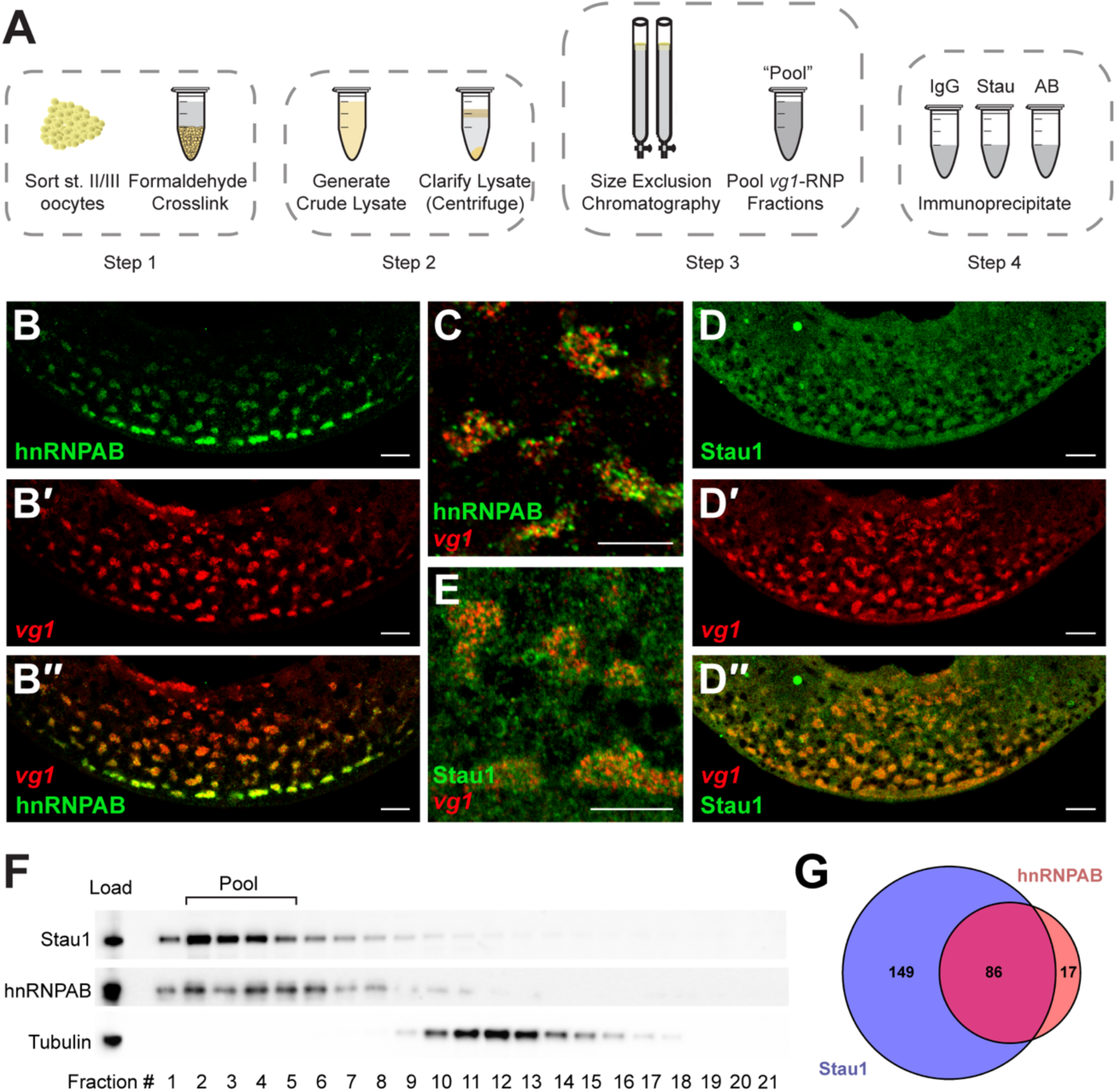
Isolation of L-bodies. **(A)** Schematic of the L-body isolation procedure. Stage II/III oocytes were formaldehyde crosslinked, homogenized, clarified by centrifugation, and fractionated by SE chromatography. Fractions containing *vg1* mRNA were pooled and divided equally for immunoprecipitations using anti-Stau1 (Stau) and anti-hnRNPAB (AB) antibodies, with IgG as a negative control. **(B)** A cryosection of a stage II oocyte is shown, with the vegetal cortex at the bottom. Combined FISH-IF was used to detect hnRNPAB protein (B, green) and *vg1* mRNA (B′, red) The merge is shown in B′′. Scale bar=20 μm. **(C)** High magnification view of the vegetal cytoplasm of a stage II oocyte showing colocalization of *vg1* mRNA (red) and hnRNPAB (green). Scale bar=10 μm. **(D)** A cryosection of a stage II oocyte is shown with the vegetal cortex at the bottom. Combined FISH-IF was used to detect Stau1 (D, green) and *vg1* mRNA (D′, red) The merge is shown in D′′. Scale bar=20 μm. **(E)** High magnification view of the vegetal cytoplasm of a stage II oocyte showing colocalization of *vg1* mRNA (red) and Stau1 protein (green). Scale bar=10 μm. **(F)** Immunoblot analysis is shown, of SE column fractions probed with anti-Stau1, anti-hnRNPAB, and anti-Tubulin. The input (load) is at the left and fraction numbers are shown at the bottom. Stau1 and hnRNPAB chromatograph primarily in the void volume; fractions 2-5, which were pooled for further purification. **(G)** Mass spectrometry was performed on parallel IPs using anti-Stau1, anti-hnRNPAB, and IgG. A Venn diagram shows the overlap between hnRNPAB (red) and Stau1 (blue); 86 proteins were identified as enriched over IgG in both the hnRNPAB and Stau1 IPs.

### L-bodies contain proteins found in other classes of cytoplasmic RNP granules

The L-body proteome (Fig. 4A) contains all known *vg1* RBPs: Elavl1, Elavl2, hnRNPAB, Vera (Igf2bp3), PTB (Ptbp1), Stau1, and Stau2 (Havin et al. 1998; Deshler et al. 1998; Cote et al. 1999; Yoon and Mowry 2004; Allison et al. 2004; Colegrove-Otero et al. 2005; Czaplinski et al. 2005; Arthur et al. 2009; Snedden et al. 2013). The presence of both Dynein (Dynll1 and Dynll2) and Kinesin (Kif3b) subunits is also notable, further suggesting the isolation of a transport competent RNP. The success in isolating the known *vg1*-interacting proteins and stringent thresholds for analyses gives a high degree of confidence that the remaining 76 proteins represent newly identified L-body components.

**Figure 4.**
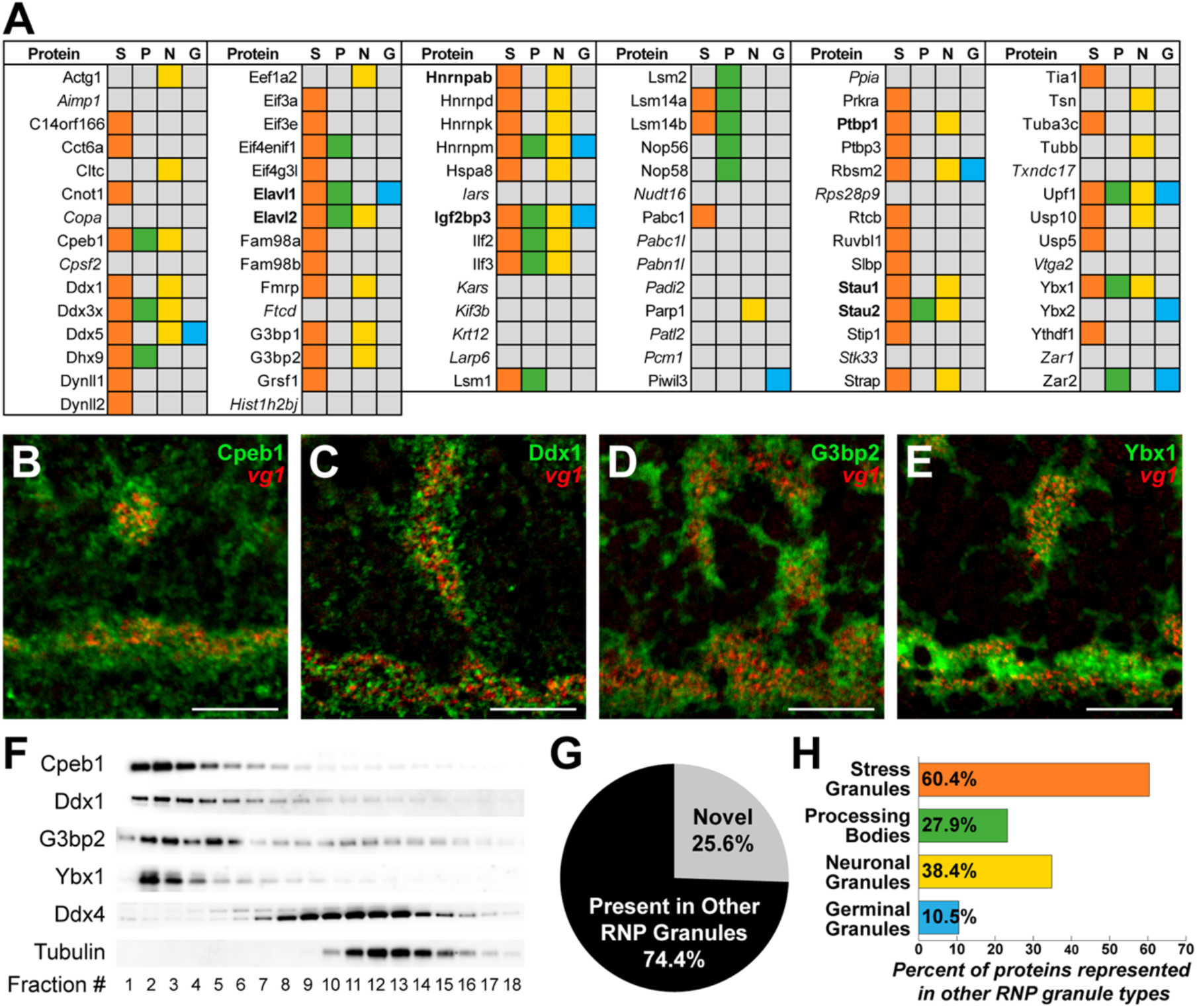
L-bodies contain protein components common to other classes of cytoplasmic RNP granules. **(A)** Protein constituents of L-bodies are also found in stress granules (S, orange), P-bodies (P, green), neuronal granules (N, yellow), and germ granules (G, blue). Proteins found in L-bodies but not reported in other granules, are italicized and previously known *vg1* RBPs are bolded. **(B-E)** Combined FISH-IF was performed to validate potential protein constituents of L-bodies. Shown are images of the vegetal cytoplasm of stage II oocytes with the vegetal cortex at the bottom; scale bars=10μm. Shown in red is *vg1* mRNA detected by FISH. Merged in green is IF using the following antibodies: **(B)** anti-Cpeb1, **(C)** anti-Ddx1, **(D)** anti-G3bp2, and **(E)** anti-Ybx1. **(F)** Immunoblot analysis is shown, of SE column fractions probed using anti-Cpeb1, anti-Ddx1, anti-Ddx3, anti-G3bp2, anti-Ybx1, anti-Ddx4, and anti-Tubulin antibodies. The fraction numbers are indicated at the bottom. **(G)** Overlap (black, 74.4%) of L-body constituents with other cytoplasmic granules; unique proteins are indicated by gray (25.6%). **(H)** The percent of identified L-body proteins that are found in other cytoplasmic RNP granule types: stress granules (60.4%, 52 of 86), P-bodies (23.3%, 24 of 86), neuronal granules (34.8%, 33 of 86), and germinal granules (10.5%, 9 of 86). See also Figures S3 and S4, and Tables S1, S2, and S3.

Putative L-body components were validated by both FISH-IF (Figs. 4B-E, S3A-J) and SE chromatography (Figs. 4F, S3K). As shown in Fig. 4, Cpeb1 (Fig. 4B), Ddx1 (Fig. 4C), G3bp2 (Fig. 4D), and Ybx1 (Fig. 4E) are each found in large *vg1* mRNA-containing structures. In addition, the candidate proteins exhibit enrichment in the vegetal cytoplasm and at the vegetal cortex (Figs. 3, S4) that is similar to the distribution of the known *vg1* RBPs hnRNPAB, Stau1, Elavl1, Elavl2, PTB, and Vera. Analysis of Cpeb1, Ddx1, G3bp2, and Ybx1 by SE chromatography shows that each protein elutes in the void volume (Fig. 4F) in large complexes that are sensitive to RNase treatment (Table S1). Ddx4, which was not identified as a potential L-body constituent by mass spectrometry (Fig. 4A), is not enriched in the void volume and is present in later fractions (Fig. 4F). Using these approaches, we validated 35 of the putative L-body components listed in Fig. 4A (Table S1, Figs. S3, S4). Gene ontology (GO) analysis (Table S2) revealed that four of the five top GO-categories are related to nucleic acid binding, with the top category being RNA binding. We also found a significant enrichment of ATP binding, in part contributed by the four DDX helicases identified in L-bodies (Fig. 4A), which may act in RNP remodeling through their ability to unwind RNA and alter RNA-protein interactions (Gilman et al. 2017).

A striking feature of the L-body proteome is the abundance of proteins that are known components of previously-described cytoplasmic RNP granules, including stress granules, P-bodies, neuronal granules, and germ granules (designated S, P, N, and G, respectively; Fig. 4A, Table S3). Remarkably, the majority (74%) of the identified L-body components are present in at least one other RNP granule type, with 38 of those (36%) being found in more than one RNP granule type (Fig. 4H). The remaining proteins (26%) appear to be novel to L-bodies (Fig. 4G) and could provide functional specificity. These results indicate that L-bodies represent a new class of cytoplasmic RNP granule.

### L-bodies exhibit non-liquid-like properties

Because many proteins found in cytoplasmic RNP granules contain intrinsically disordered regions (IDRs; Uversky 2017), we also examined the prevalence of IDRs in the L-body proteome. Using SLIDER (Peng et al. 2014), a predictive tool for IDRs, we found the majority (74%) of all L-body components contain a putative IDR, which is a significant enrichment over the *Xenopus* proteome (56%; Fig. 5A). Other types of cytoplasmic RNP granules have been described as phase-separated bodies, with constituent proteins and mRNAs exhibiting liquid-like behavior in some RNP granules and gel- or amyloid-like properties in others (reviewed in Banani et al. 2017; Shin and Brangwynne 2017). Intriguingly, prion-like domains, which are a specialized category of IDRs that are overrepresented in gel- and amyloid-like structures, are also overrepresented in the L-body proteome (Fig. S5). To probe the properties of L-bodies, we first stained oocyte sections with thioflavin, a dye that exhibits a selective fluorescence shift in the presence of cross-beta strands, which are characteristic of amyloid structures (Guntern et al. 1992). The vegetal oocyte cytoplasm stains richly with thioflavin (Fig. 5B), revealing a mesh-like substructure evident at high magnification (Fig. 5C). To test whether the thioflavin-staining structures correspond to L-bodies, we next used combined FISH and thioflavin staining to ask whether vegetally-localized RNAs co-localize with the mesh-like structures. Remarkably, localized mRNAs, *vg1* (Fig. 5D), *vegT* (Fig. 5E), and *trim36* (Fig. 5F) are highly coincident with the thioflavin staining. By contrast, non-localized *gapdh* mRNA is not co-localized with the thioflavin staining (Fig. 5G). The distinct staining with thioflavin, revealing an apparent mesh-like structure, suggests that L-bodies may not fall into the class of liquid-like granules and may instead display gel- or solid-like properties. To test this idea, we prepared lysates from oocytes that were injected with fluorescent *LE* RNA (as in Fig. 2A,B,E) to mark the L-bodies. After SE chromatography (as in Fig. 3F), the “*ex vivo*” L-bodies were treated with 1,6-hexanediol, which disrupts liquid-like granules (Kroschwald et al. 2015). The *ex vivo* L-bodies, which are evident as fluorescent spheres (Fig. 5H), are insensitive to this treatment (Fig. 5 H-I), suggesting that *ex vivo* L-bodies are not liquid-like in nature. These results raise the possibility that L-bodies may instead have more gel- or solid-like properties.

**Figure 5.**
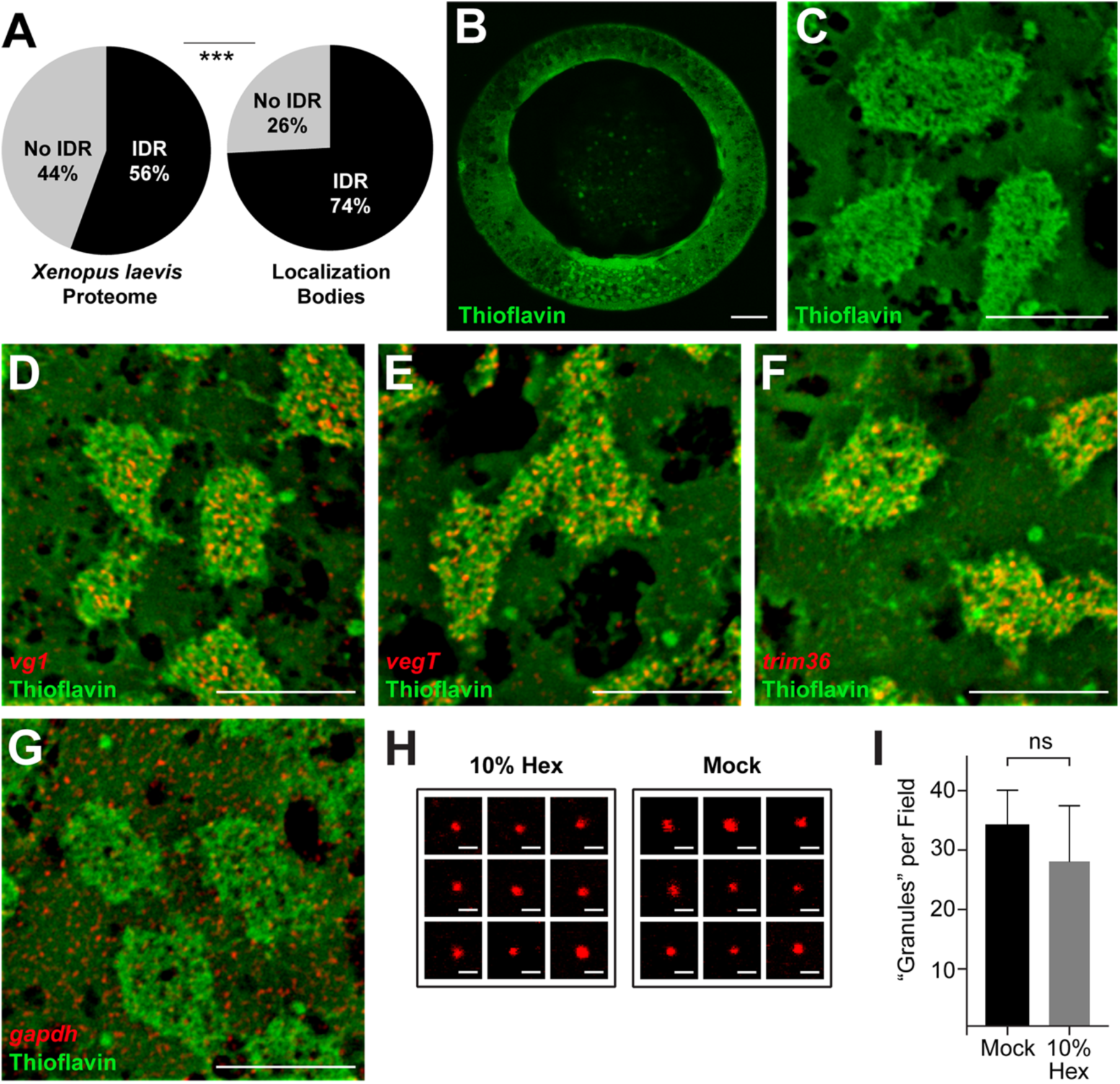
L-bodies exhibit a mesh-like structure and non-dynamic characteristics. **(A)** Comparison by SLIDER analysis (Peng et al. 2014) for long stretches (>30 consecutively disordered residues) IDRs for the *X. laevis* proteome *vs*. the L-body proteome. *** indicates p < 0.001. See also, Figure S5. **(B)** A stage II oocyte stained with thioflavin is shown, with the vegetal cortex at the bottom. Scale bar=50 μm. **(C)** High magnification view of the vegetal cytoplasm of a stage II oocyte stained with thioflavin. Scale bar=10 μm. **(D-G)** High magnification views of the vegetal cytoplasm of stage II oocytes stained with thioflavin are shown; scale bars=10 μm. Thioflavin staining was combined with FISH for detection of the following RNAs: **(D)** *vg1* RNA, **(E)** *vegT* RNA, **(F)** *trim36* RNA, **(G)** *gapdh* RNA. **(H)** L-bodies labeled with fluorescent *LE* RNA were enriched by SE chromatography and treated for 15 minutes at 18°C with 10% 1,6-hexanediol (10% Hex, left) or “Mock” control (right). Scale bars=2μm. **(I)** The number of fluorescent granules was quantitated after treatment with 10% 1,6-hexanediol (gray) or mock treatment (black). Error bars indicate standard error of the mean. See also Figure S6.

### Dynamics of L-body RNAs and proteins differ *in vivo*

To examine the dynamics of L-bodies *in vivo*, protein mobility was assessed by fluorescence recovery after photobleaching (FRAP) in live oocytes. In these experiments, we marked L-bodies by microinjection of fluorescently-labeled *LE* RNA into stage II oocytes along with mRNA expressing mCherry-tagged L-body proteins (Fig. 6A-B). To determine dynamics within the L-bodies, partial granule FRAP was performed on three previously known L-body proteins (hnRNPAB, Stau1 and Vera) as well as a newly-identified constituent, Ybx1 (Tafuri and Wolffe 1990). Each of these proteins contain at least one RNA-binding domain and both hnRNPAB and Ybx1 additionally contain IDRs. Each of the four RBPs exhibited significant mobility within the L-body, with hnRNPAB showing the highest mobile fraction (94.2%) and Vera having the lowest (52.9%; Fig. 6A). These data indicate that protein constituents of L-bodies are dynamic, which was unexpected due to the thioflavin staining and the 1,6-hexanediol insensitivity (Fig. 5). These results suggest that L-bodies may be multiphase condensates, with a subclass of L-body constituents exhibiting dynamic properties and others being more gel- or solid-like.

**Figure 6.**
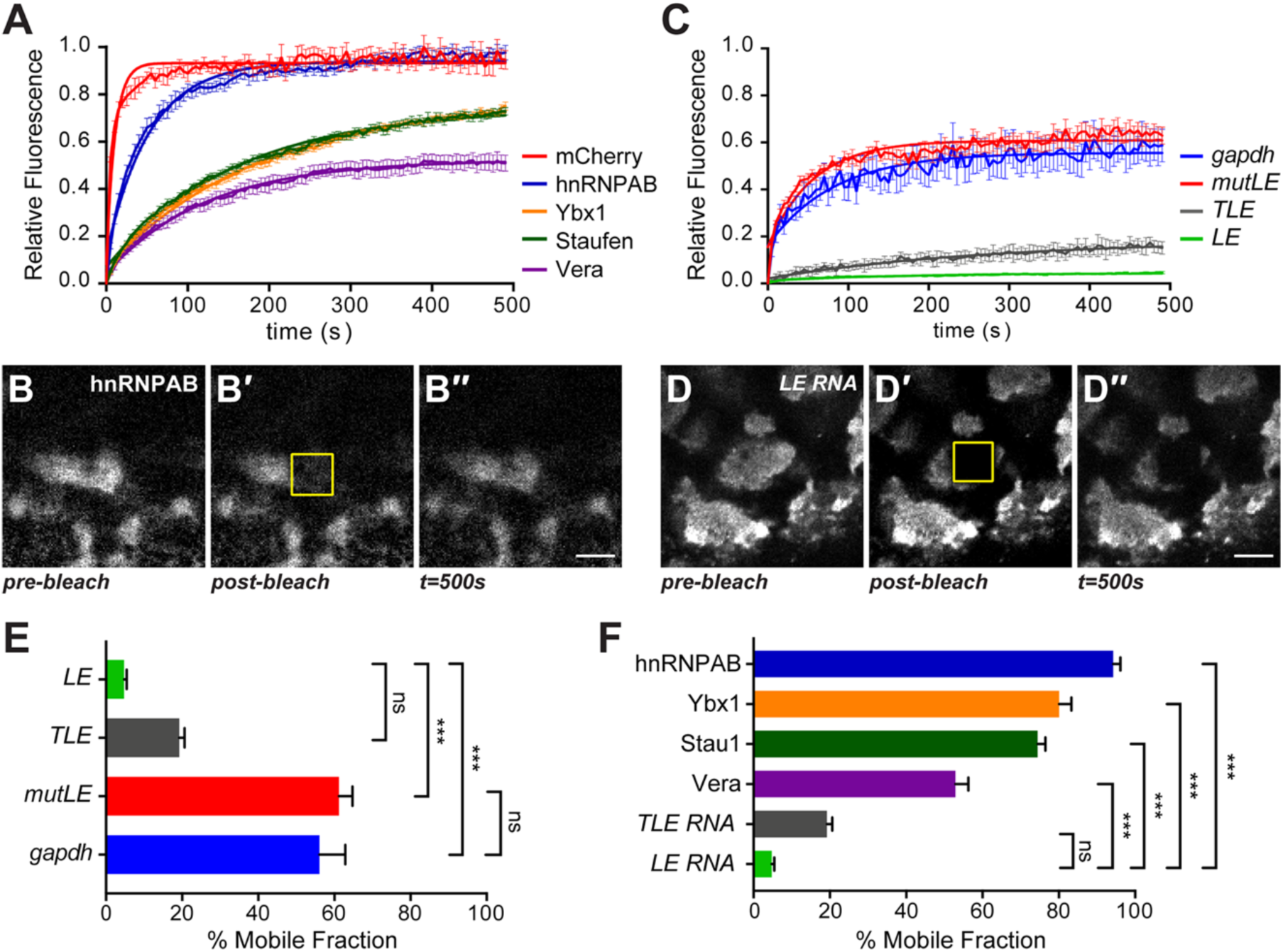
L-bodies contain dynamic proteins and non-dynamic RNAs. **(A)** Stage II oocytes expressing mCherry (mCh), or the following mCh-tagged proteins: hnRNPAB, Stau1, Ybx1, Vera, were microinjected with Cy5-labeled *vg1 LE* RNA to mark L-bodies. Recovery curves are shown for mCherry (red), mCh-hnRNPAB (blue), Stau1-mCh (green), Ybx1-mCh (orange), and Vera-mCh (purple). Error bars indicate standard error of the mean. **(B)** An image of the vegetal cytoplasm of an oocyte microinjected with mCh-hnRNPAB is shown, with a 10 μm^2^ ROI (yellow); scale bar=10 μm. B′ and B′′ show the post-bleach and 500 second (s) timepoints, respectively. Photobleaching was corrected using the ImageJ plugin CorrectBleach V2.0.2. Zenodo (Miura et al., 2014) **(C)** Stage II oocytes were microinjected with Cy3-labeled *vg1 LE* RNA to mark L-bodies and co-injected with the following Cy5-labeled RNAs for FRAP analysis: *vg1 LE*, mutated *vg1 LE* (*mutLE*), *vegT LE* (*TLE*), or *gapdh*. Recovery curves are shown for *LE* RNA (green), *mutLE* RNA (red), *TLE* RNA (gray), and *gapdh* RNA (blue). Labeling of RNAs with Cy3 *vs*. Cy5 does not affect dynamics (see Fig. S7). Error bars show standard error of the mean. **(D)** An image of the vegetal cytoplasm of an oocyte microinjected with Cy5-labeled *LE* RNA is shown, with a 10 μm^2^ ROI (yellow); scale bar=10 μm. D′ and D′′ show the post-bleach and 500 second (s) timepoints, respectively. Photobleaching was corrected using the ImageJ plugin CorrectBleach V2.0.2. Zenodo (Miura et al., 2014) **(F)** The mobile fraction is shown for *LE* RNA (green, 4.8%±0.6%), *TLE* RNA (gray, 19.2±1.4%), *mutLE* RNA (red, 61.2%±3.6%), and *gapdh* RNA (blue, 56.1±6.7%), determined by FRAP as in panel C. The results from 7 oocytes per RNA are shown and error bars show standard error of the mean. P-values were calculated using one-way ANOVA with Tukey’s multiple comparison correction; ***p<0.001, ns is not significant. **(F)** The mobile fraction for mCh-hnRNPAB (blue, 94.2±1.9%), Stau1-mCh (dark green, 74.5±2.0%), Ybx1-mCh (orange, 80.1±3.2%), Vera-mCh (purple, 52.9±3.4%), *TLE* RNA (gray, 19.2±1.4%), and *LE* RNA (green, 4.8%±0.6%) was determined by FRAP, as in panels A and C. The results from 7 oocytes per protein and RNA are shown and error bars indicate standard error of the mean. P-values were calculated using one-way ANOVA with Tukey’s multiple comparison correction; ***p<0.001, ns is not significant. See also Table S4.

The distinct thioflavin staining of L-bodies (Fig. 5 B-G) suggests that they may contain one or more non-dynamic constituents, in addition to the dynamic protein constituents (Fig. 6A). We first examined the distribution of L-body proteins by IF, and none of the 46 proteins tested exhibited a mesh-like pattern similar to the thioflavin staining pattern (Fig.S3 and data not shown). Because thioflavin may also stain structured RNAs (Xu et al. 2016), we tested whether *Xenopus* oocyte RNA is capable of staining with thioflavin. Indeed, *Xenopus* oocyte RNA, which forms droplets at low concentrations and gels at high concentrations (Fig. S6B,C), binds to thioflavin in a time- and concentration-dependent manner (Fig. S6A). To examine the dynamics of L-body RNAs *in vivo*, we performed FRAP on microinjected RNAs. Both *vg1 LE* (*LE*) and *vegT LE* (*TLE*) RNAs are highly immobile *in vivo*, with mobile fractions of 4% and 19%, respectively (Fig. 6C-E). In contrast, non-localized RNAs are much more dynamic; the mobile fractions for *mutLE* RNA and *gapdh* mRNA are 61.2% and 56.1%, respectively (Fig. 6D,F). Notably, mobility does not correlate with length of either the proteins or the RNAs (Table S4). The low mobility of localized RNAs compared with the dynamic behavior of constituent proteins (Fig. 6F) suggests at least two distinct phases within L-bodies, with RNA as a potentially solid- or gel-like phase enveloped in a dynamic protein phase.

## Discussion

In this work, we have discovered that vegetal mRNA localization in *Xenopus* oocytes proceeds via the formation of L-bodies, which are a new class of cytoplasmic RNP granules. We base this conclusion on several lines of evidence: First, L-bodies are large cytoplasmic RNPs that are specifically enriched for vegetally localized mRNAs (Figs. 1-2). Second, incorporation into L-bodies is determined by *cis*-elements within mRNAs and enrichment is required for subsequent localization. Third, the protein composition of purified L-bodies exhibits a high degree of similarity to that of other classes of cytoplasmic RNP granules (Figures 3-4). In addition to over two-thirds of the proteins being directly conserved amongst cytoplasmic RNP granules, the over-representation of multivalent RBPs and IDR-containing proteins (Fig. 5) provides a compositional link between L-bodies and other classes of phase separated RNPs (reviewed in Wang et al. 2014; Banani et al. 2017; Shin and Brangwynne 2017; Putnam et al. 2019). Finally, we find that L-bodies are multi-phase RNP granules composed of dynamic proteins and non-dynamic localized RNAs (Fig. 6), with RNA potentially serving as a scaffold component.

Maternal mRNAs are transported within developing oocytes of *Xenopus* and many other species where their local translation is critical for proper embryonic patterning. Motor-based transport of such mRNAs relies on the assembly of RNP transport cargos, a conserved feature of mRNA localization pathways (Gagnon and Mowry 2011). Yet, the molecular and physical nature of these cargos has remained largely unknown. Our work establishes that transport cargos in *Xenopus* oocytes are large cytoplasmic RNP granules. We find that L-bodies, which contain many copies of localized mRNAs, are surrounded by a dense network of microtubules that are decorated by molecular motors with known roles in vegetal mRNA transport (Betley et al. 2004; Messitt et al. 2008; Gagnon et al. 2013) (Fig. 1). In contrast with some other types of transport cargos, where mRNAs are packaged singly (Batish et al. 2012; Buxbaum et al. 2014; Little et al. 2015), multiple localized transcripts are contained within each L-body (Fig. 1). Importantly, vegetally-localized RNA sequences are specifically incorporated into L-bodies, while non-localizing RNAs are not, regardless of length (Figs. 1-2), suggesting that enrichment in L-bodies is required for vegetal mRNA localization. New insight into the molecular and physical nature of these transport cargos came from our purification and mass spectrometry results, which revealed that L-bodies contain protein constituents that are shared among many cytoplasmic RNP granules (Figs. 3-4). Taken together, these results show that vegetal mRNAs are transported in L-bodies, which are a new type of cytoplasmic RNP.

A clear substructure of L-bodies is apparent upon staining with thioflavin (Fig. 5), a dye that typically stains amyloid-like aggregates formed by protein β-sheet interactions (Nilsson 2004). While extensive efforts to identify protein components with both structured patterns of L-body localization and non-dynamic intra-granule motility did not provide insights into the composition of the mesh-like substructure, evidence of a role for RNA emerged. The overlap of thioflavin staining with localized mRNA (Fig. 5D-F) raises the possibility that the mesh-like substructure may be composed of RNA. This is particularly intriguing given recently observed functions for mRNA structure in liquid-liquid phase separation and transitions to gel- or solid-like phases (Jain and Vale 2017; Langdon et al. 2018). RNA-RNA interactions have also been implicated in RNA transport and localization. In *Drosophila* oocytes homotypic intermolecular RNA-RNA interactions are necessary for localization of *bicoid* and *oskar* mRNAs (Ferrandon et al. 1997; Jambor et al. 2011). In *Xenopus* oocytes, depletion of *vegT* mRNA, a L-body component, leads to the mislocalization of *vg1* mRNA (Heasman et al. 2001), suggesting a role for heterotypic mRNA interactions. Notably, thioflavin has been demonstrated, both here (Fig. S6) and elsewhere (Xu et al. 2016), to recognize dense networks of RNA structure, suggesting that RNA may be the source of thioflavin staining observed in L-bodies.

Our results using *in vivo* imaging strategies and *ex vivo* analysis of purified L-bodies to probe their physical nature reveals that L-bodies are multiphase condensates containing both nondynamic RNAs and dynamic proteins. With RNA as a potential scaffold, dynamic client proteins are both enriched over and actively exchanging with the surrounding cytoplasm (Figs. 3-6). This multiphase body composition parallels findings in P granules (Wang et al. 2014; Putnam et al. 2019) and stress granules (Jain et al. 2016) and may reflect a fundamental aspect of RNP granule organization. However, unlike in other examples, our findings suggest that in L-bodies, RNA is a critical scaffold component, as localized RNAs are non-dynamic within L-bodies (Fig. 6). While RNA has been viewed as a multivalent platform, capable of mediating intermolecular interactions between *trans*-acting proteins, recent studies have challenged this ancillary role for RNA in granule organization, and offered new roles for RNA-RNA interactions in defining the composition of RNP granules (Jain and Vale 2017; Van Treeck et al. 2018; Langdon et al. 2018; Niepielko et al. 2018; Yamazaki et al. 2018). Interestingly, within the dynamic phase of L-bodies, protein mobility may correlate with the degree of resident valency, as Vera which harbors multiple sequence-specific RNA-binding domains shows reduced mobility relative to lower valency RBPs (Fig. 6). These data support a model in which RNP granule composition is specified by heterotypic multivalent interactions (Banani et al. 2016), wherein the intra-granular behavior of a client protein is dictated by its potential to interact with multivalent RNA scaffolding.

The extremely low mobility of localized RNAs within L-bodies further suggests an important distinction between the localizing RNA containing portion of this RNP granule and non-localizing RNAs. While non-localized mRNAs are not concentrated within the L-body, they are not excluded (Figs. 1-2). Interestingly, non-localizing mRNAs, including the length-matched mutant *vg1 LE* (*mut-LE*) and *gapdh*, may also remain in dynamic exchange with the surrounding cytoplasm (Fig. 6). Incorporation into L-bodies relies on sequence-encoded features of the RNAs (Fig. 2; Lewis et al. 2008) and likely represents a capture event of freely diffusing RNA. This is supported by the observation that exogenously expressed transcripts are recruited into preexisting L-bodies, rather than packaged independently (Fig. 2). Importantly, the observation that sequence mutations within the *vg1 LE* that are known to ablate localization (Lewis et al. 2008) also block L-body enrichment (Fig. 2), functionally links L-bodies to mRNA localization.

We propose a model for L-body structure and assembly (Fig. 7). Localized RNAs provide an organizational scaffold, wherein interactions with specific RBPs both promotes the local enrichment of L-body factors and controls specific incorporation of localized mRNAs into L-bodies. As localized RNAs become strongly enriched, high RNA concentrations promote RNA-RNA interactions and the formation of a solid- or gel-like RNA phase. Thus, RNA acts as a scaffold and self-assembly drives the observed multiphase behavior of L-bodies. Protein constituents become further enriched in L-bodies through their interaction with the RNA scaffold, while non-localized RNAs and other proteins, which lack the capacity to interact with the RNA scaffold, freely diffuse in and out of L-bodies.

**Figure 7.**
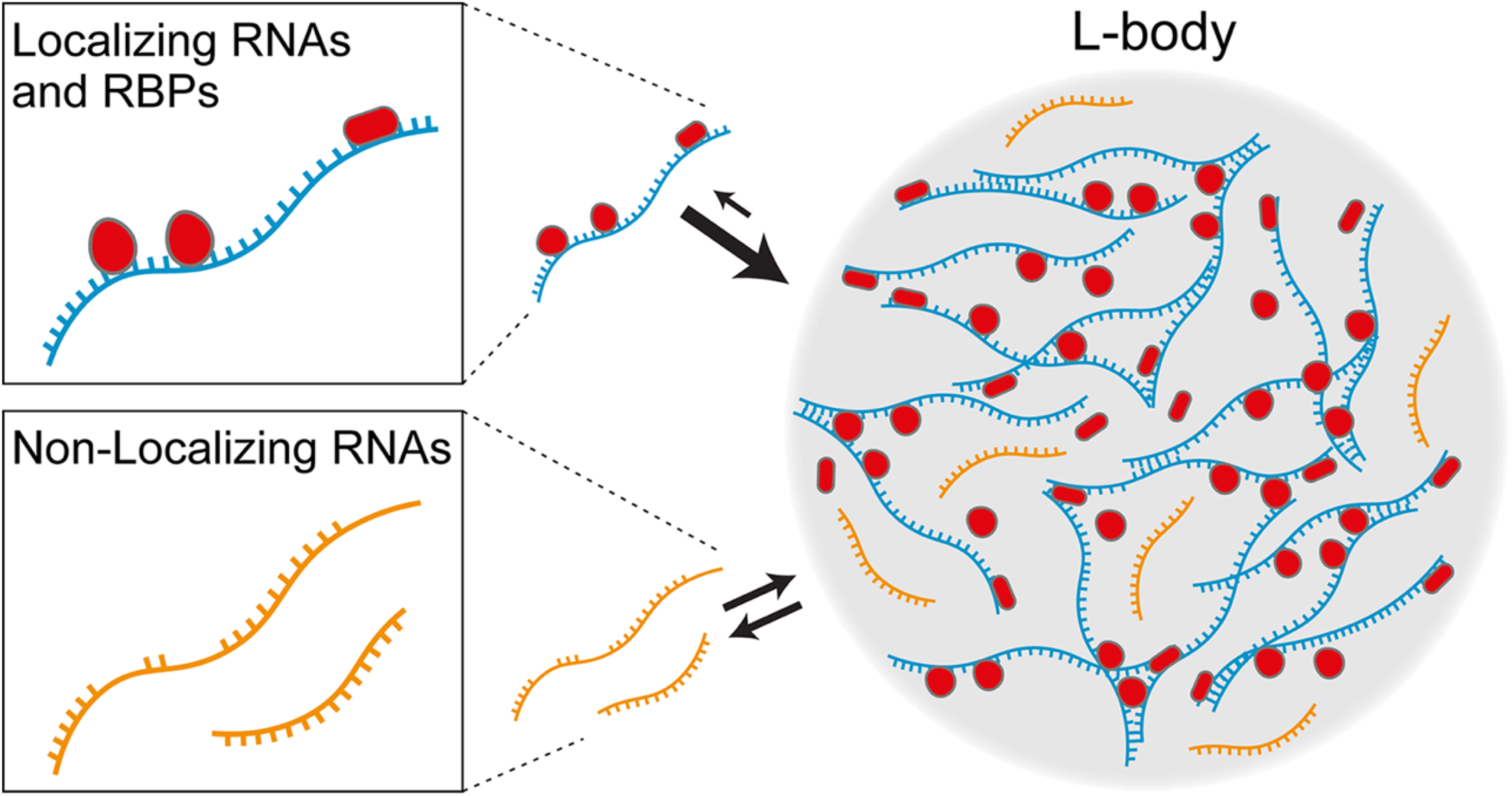
Model for L-body structure and assembly. Localizing RNAs (blue) interact with specific RNA-binding proteins (RBPs) that facilitate local enrichment. Non-localizing RNAs (gold) lack binding sites for these RBPs and do not condense with other localized RNAs. Specific RBPs (red) facilitate enrichment (large arrows) of localized RNAs in L-bodies (gray). L-body enrichment results in high local RNA concentrations, facilitating RNA-RNA interactions and formation of a gel-like phase. Non-localized RNAs (and proteins) are not excluded from L-bodies, and can freely move in and out (double arrows).

By defining the physical cargo of vegetal transport in *Xenopus* oocytes as multiphase RNP granules (L-bodies), regulation of distinct transport and developmental steps can be understood in the context of the unique physical properties of phase-separated structures. Cytoplasmic RNP granule components have been observed to transition between diffuse, liquid-like, or more solid states in response to environmental cues that regulate coassembly, viscosity, and demixing specificity within granules (Hubstenberger et al. 2013; Elbaum-Garfinkle et al. 2015; Hubstenberger et al. 2015; Jain et al. 2016; Putnam et al. 2019). In this way, the innately reversible interactions that underlie RNP granule composition allow for fine-tuning of RNP granule dynamics, which can be exploited as a regulatory mechanism. In oocytes, which are non-cycling for protracted periods of time, packaging of maternal mRNAs into non-dynamic gel- or solid-like RNP granules, such as L-bodies, may be an important mechanism to silence translation over long timeframes. Upon response to appropriate developmental cues, these stable RNP granules must then be disassembled. The mechanisms regulating stabilization and disassembly in oocytes are likely to represent general paradigm for understanding regulation of dynamic phase transitions in space and time.

## Materials and Methods

### Oocyte Isolation and Culture

Oocytes were harvested from *Xenopus laevis* females, either J Strain (NXR, catalog # NXR_0024) or wild type (Nasco, catalog # LM00535MX). All animal experiments were approved by the Brown University Institutional Animal Care and Use Committee. Oocytes were enzymatically defolliculated in 3 mg/ml collagenase (in 0.1 M KPO_3_) followed by washes in MBSH (88mM NaCl, 1mM KCl, 2.4mM NaHCO_3_, 0.82mM MgSO_4_, 0.33mM CaCl_2_, 0.33mM Ca(NO_3_)_2_, 10mM HEPES pH7.6). Stage II-III oocytes were cultured at 18°C in XOCM [50% Leibovitz L-15, 15mM HEPES (pH 7.6), 1mg/mL insulin, 50U/mL nystatin, 100U/mL penicillin/streptomycin, 0.1mg/mL gentamicin].

### RNA Transcription and Microinjection

Fluorescently-labeled RNAs, *vg1 LE* (Gautreau et al. 1997), *vegT LE* (*TLE*) (Bubunenko et al. 2002), nonlocalizing mutant *vg1 LE* (*mutLE*) (Lewis et al. 2008), and *βglobin* (Krieg and Melton 1984) were generated from linearized plasmids pSP73-2×135, pSP73-VegTLE, pSP73-VLEΔE2ΔVM1, and pSP64-XBM respectively. For transcription of *gapdh* a PCR product was generated using gapdh-specific primers (gapdh T7 Fwd and gapdh Rev), with the forward primer containing the T7 promoter sequence. Transcription was performed using the MEGAscript T7 transcription kit in the presence of 250 nM Cy™ 3- or Cy™ 5-UTP. mCherry (mCh)-tagged RNAs were generated using the mMessage machine transcription kit from linearized plasmids, as follows: pSP64:mCh-hnRNPABx2, pSP64:Stau1-mCh, pSP64:Vera-mCh, and pSP64:YBX1-mCh. Barcoded RNAs were generated from PCR products using pSP73-2×135, pSP73-VLEΔE2ΔVM1, and pSP64-XBM plasmids as templates and the following primers: LE barcode A Fwd, mutLE barcode B Fwd, LE/mutLE Rev, Bglobin barcode A Fwd, Bglobin barcode B Fwd, Bglobin barcode C Fwd, Bglobin Rev. Cy-labeled RNAs were injected at 200 nM, mCh RNAs were injected at 500 nM and barcoded RNAs were injected at 1.25 nM. Following microinjection, oocytes were cultured for 16-24 hrs. at 18°C in XOCM.

### Fluorescence in situ hybridization and immunofluorescence

Oocytes were fixed for 1 hour at 22°C in FTG (80 mM K PIPES, pH6.8, 1 mM MgCl_2_, 5 mM EGTA, 0.2% Triton X-100, 3.7% formaldehyde, 0.25% glutaraldehyde, 0.5 μm paclitaxel), followed by post-fixation in 100% methanol overnight at 22°C (Becker and Gard 2006). After re-hydration in TBS, oocytes were incubated overnight at 4°C in 100 mM NaBH_4_ in TBS (Becker and Gard 2006). After 2 washes in TBSN (0.1% NP-40 in TBS), followed by 2 washes in TBSNB (0.2% ultrapure BSA in TBSN), oocytes were equilibrated stepwise in TBSNB-30S (30% sucrose in TBSNB) and, after equilibration in an embedding mold, TBSNB-30S was replaced with OCT compound and snap frozen in an ethanol-dry ice bath.

Cryosectioning and fluorescence in situ hybridization (FISH) was performed as described in Neil and Mowry (2018), with custom FISH probes for *gapdh, trim26, vegT*, and *vg1*, applied at 500 nM. To assess incorporation of injected *LE* RNA into endogenous L-bodies, a modified *vg1*δLE probe set was engineered to exclude probes from the *vg1 LE* sequence. For combined FISH-IF, slides were subsequently washed twice in TBSNB, incubated for 3 hours at 22°C in TBS-plus (5% v/v normal goat serum, 2 mg/ml ultrapure BSA in TBSN), and incubated overnight at 4°C with primary antibodies at 20 μg/ml in TBS-plus. Primary antibodies were directly labeled with Alexa-405, Alexa-546, or Alexa-647 using Zenon labeling kits. After four 10-minute washes in TBSNB at 22°C, slides were mounted with ProLong Gold Antifade. For thioflavin staining, cryosections were incubated in thioflavin S solution (1% in 80% ethanol) for 20 min, at room temperature, followed by washes with 80% ethanol, 70% ethanol, and H_2_O. Slides were either mounted in Prolong Gold Antifade or processed for FISH as described previously (Neil and Mowry 2018) with probe sets restricted to Quasar 670 to avoid overlap with the thioflavin fluorescence. Imaging was performed on a Zeiss LSM 800 Confocal Laser Scanning Microscope.

### Fluorescence Recovery After Photobleaching (FRAP)

Two nl of RNA encoding mCh-tagged proteins (at 500 nM) or fluorescently-labeled RNAs (at 200nM) were injected into stage II oocytes and allowed to localize for 48 hours in XOCM for 18°C. Imaging was performed on an Olympus FV3000 confocal microscope with a UPLSAPO 30× oil objective (1.05NA). A 10μm^2^ ROI was bleached using a 561nm laser at 100% for 2 seconds. Recovery was monitored at five second intervals for 100 iterations. FRAP calculations were as previously described (Gagnon et al. 2013) and curve fitting was performed in GraphPad Prism 8 using Y=Y0 + (Plateau-Y0)*(1-exp(-K*x).

### Oocyte lysate preparation

Approximately 2500 stage II-III oocytes were washed with PBS, followed by crosslinking (0.1% formaldehyde in PBS) for 10 minutes at 22°C. The reaction was quenched with 0.25 M glycine (in 25 mM Tris pH 7.4,) for 5 minutes at 22°C. Oocytes were washed in <u>c</u>olumn <u>r</u>unning <u>b</u>uffer (CRB; 0.05% NP-40, 1 mM DTT, 10 mM Hepes 7.4, 100 mM KOAc, 3 mM MgOAc, 5 mM EGTA, 100 mM sucrose, 2 nU/ml Ribolock RNase Inhibitor, 1× HALT Protease Inhibitors) and homogenized in CRB (∼20 oocytes/μl). The lysate was clarified by centrifugation at 10,000×*g* for 10 minutes at 4°C and adjusted to a final protein concentration of 50 mg/ml in CRB.

### Size Exclusion Chromatography

Oocyte lysate was chromatographed on Sephacryl S400 HR resin in CRB. Column fractions were analyzed by qRT-PCR for *vg1* mRNA and by immunoblot for proteins of interest. Fractions containing the *vg1* RNA peak were pooled for further analysis. For *ex vivo* analyses, oocytes were microinjected with 2nl of 1μM Cy-labeled *LE* RNA prior to oocyte lysate preparation and chromatography. For RNase treatment of oocyte lysates prior to S400 chromatography, Ribolock RNase Inhibitor was omitted from all buffers, and RNase A was added to a final concentration of 0.1 μg/μl and incubated for 10 min. at 37°C. To control for RNA extraction efficiency, 2.5 pg of *luciferase* RNA was added to each column fraction. RNA was isolated using the Direct-Zol RNA MicroPrep kit, and cDNA was prepared from 10 μl of RNA using the iScript cDNA synthesis kit. For analysis of *vg1* mRNA distribution across fractions, qRT-PCR was performed using Luna One-Step qPCR with forward (qPCR vg1 Fwd) and reverse (qPCR vg1 Rev) *vg1* primers, and forward (qPCR luc Fwd) and reverse (qPCR luc Rev) *luciferase* primers. Copy numbers of *vg1* and *luciferase* RNAs were determined by standard curve and *vg1* copy number was divided by *luciferase* copy number per fraction. For analysis of injected RNA distribution across column fractions, qRT-PCR was performed using Luna One-Step qPCR using the following primers: Barcode A qPCR Fwd, Barcode B qPCR Fwd, Barcode D qPCR Fwd, Barcode C qPCR Rev.

### Immunoprecipitation and Immunoblotting

Antibodies were covalently crosslinked to Dynabeads, which were washed in 0.1% BSA for 5 minutes at 22°C, blocked for 1-hour in RNA blocking solution (0.05% CHAPS, 0.32 μg/μl torula RNA, PBS), and washed three times in CRB. S400 *vg1* RNA peak “pool” was incubated for 6 hours at 4°C with antibody-coupled Dynabeads in PIP buffer (50 mM Tris pH 7.4, 0.1% NP40, 0.05 mM MgCl_2_, 2 nU/ml Ribolock RNase Inhibitor, and 1× HALT Protease Inhibitors) for protein co-IP, or in RIP buffer (25 mM Tris pH 7.4, 0.5% NP40, 0.5 mM DTT, 150 mM KCl, 5 mM EDTA, 2 nU/ml Ribolock RNase Inhibitor, 1× HALT Protease Inhibitors) for RNA co-IP. For RNA-IPs, two step qRT-PCR was performed using SybrGreen Powerup MasterMix. For mass spectrometry, bound proteins were eluted with 50 mM Tris pH 8.0, 0.2% SDS, 0.1% Tween-20 at 25°C, followed by incubation at 70°C for 1 hour to reverse the crosslinking and removal of Tween-20 using HiPPR columns. For immunoblotting, antibodies were used at 1:1000, except for PTB (1:2000), Stau1 (1:2000), Tubulin (1:500), Vera (1:2000), and Ybx1 (1:100). Secondary antibodies (goat anti-rabbit IgG or goat anti-mouse IgG) were used at 1:10,000.

### Mass Spectrometry

LC-MS/MS was performed at the COBRE CCRD Proteomics Core Facility (RI Hospital) on an automated platform (Yu and Salomon 2009, 2010) connected to a Q Exactive Plus mass spectrometer (Thermo Fisher Scientific) as detailed in (Ahsan et al. 2017). Peptide spectrum matching of MS/MS spectra of each file was searched against a species-specific database (*Xenopus laevis;* <u>UniProt</u>; downloaded 2/1/2015) using MASCOT v. 2.4. A concatenated database containing “target” and “decoy” sequences was employed to estimate the false discovery rate (FDR) (Elias and Gygi 2007). Msconvert v. 3.0.5047, using default parameters and with the MS2Deisotope filter on, was employed to create peak lists for Mascot. The Mascot database search was performed with the following parameters: trypsin enzyme cleavage specificity, 2 possible missed cleavages, 10 ppm mass tolerance for precursor ions, 20 mmu mass tolerance for fragment ions. Search parameters permitted variable modification of methionine oxidation (+15.9949 Da) and static modification of carbamidomethylation (+57.0215 Da) on cysteine. The resulting peptide spectrum matches (PSMs) were reduced to sets of unique PSMs by eliminating lower scoring duplicates. To provide high confidence, the Mascot results were filtered for Mowse Score (>20). Peptide assignments from the database search were filtered down to a 1% FDR by a logistic spectral score as previously described (Elias and Gygi 2007; Yu et al. 2009). Mass spectrometry proteomics data have been deposited to the ProteomeXchange Consortium via the PRIDE via the PRIDE (Perez-Riverol et al. 2019) partner repository with the dataset identifier PXD013742.

### Gene Ontology, and Domain Analysis

Gene Ontology (GO) analysis was determined using DAVID (Huang et al. 2009b, 2009a), with *Xenopus tropicalis* as background. Identification of putative IDRs were determined using SLIDER (Peng et al. 2014), which predicts long disordered segments (>30 consecutive disordered residues), against the hnRNPAB/Stau1 dataset or the *Xenopus laevis* J-strain 9.2 genome (xenbase.org). Only those proteins with a SLIDER score greater than 0.55 were considered likely to have an IDR. Prion-like domains were calculated using PLAAC (Lancaster et al. 2014) with default settings. Only those proteins with a COREscore greater than zero were considered to have a prion-like domain.

### Quantification and Statistical Analysis

For mass spectrometry bioinformatics and analysis, peptide counts were generated by the Rhode Island Hospital, COBRE CCRD Proteomics Core Facility affiliated with Brown University. Downstream analysis was performed in-house using R/R Studio (v3.5.0/v1.1.447 & v1.1.453). For Stau1 and hnRNPAB IPs, only those proteins with peptides with at least 3 replicates were considered. Enrichment over IgG was determined if the sum of peptides across all four biological replicates was as least two-fold greater than the sum of the peptides in all four IgG experiments. For unannotated protein hits (i.e.: LOC, MGC, Xelaev, Xetrov delimitators), peptides were BLAST against the *Xenopus laevis* JGI 9.1v1.8.3.2 annotation database, available from xenbase.org. Area proportional venn diagrams were produced using the VennDiagram package for R (v3.5.0).

All bioinformatics analysis was carried out with either R/R Studio (v3.5.0/v1.1.447 or v1.1.453) or Microsoft Excel (Office 365) software. For RNA-IP analysis, LinRegPCR (Ramakers et al. 2003) was used to calculate primer efficiency for both *vg1* and *luciferase* primers. The geometric mean of primer efficiency was calculated in Excel for each primer across four experimental replicates. The relative fold enrichment was calculated using the Pfaffl method (Pfaffl 2001), and the geometric mean efficiency of *vg1* and *luciferase* primers as determined by LinRegPCR (Ramakers et al. 2003). Log2 fold enrichment of *vg1* in RBP-IPs was calculated relative to IgG and normalized to *luciferase*. Four replicates were performed, and a Mann-Whitney U test was performed for statistical significance (De Neve et al. 2013) using R studio. Statistics relating to relative enrichment of gene ontology annotations, intrinsically disordered domains, and prion-like domains were performed in R/R studio (v3.5.0/v1.1.453) using the Fisher’s exact test.

### Data and Software Availability

The *ex vivo* RNP granule analysis script was developed in-house and is freely available as an ImageJ plugin at: https://github.com/sjeschonek/mowrylab_rna_localization/blob/master/ExVivo_GranuleAnalysis.py.

## Acknowledgments

We thank J. Fallon, N. Fawzi, E. Larschan, and J. Otis for comments on the manuscript. This work was funded by grant R01-GM071049 from the NIH to KLM. CRN, SPJ, and LCO were supported in part by NIH grant T32-GM07601. Mass spectrometry was performed in the Rhode Island Hospital, COBRE CCRD Proteomics Core Facility (supported by P30-GM110759) with advice and technical support from A.R. Salomon and N. Ahsan.

## Author Contributions

CRN, SPJ, SEC, LCO, EAP, TAW, and KLM designed, carried out, and/or analyzed experiments. CRN, SPJ, and KLM drafted the manuscript. CRN, SPJ, SEC, LCO, EAP, and KLM contributed to review and editing of the manuscript.

## Supplemental Figure Legends

**Figure S1.**
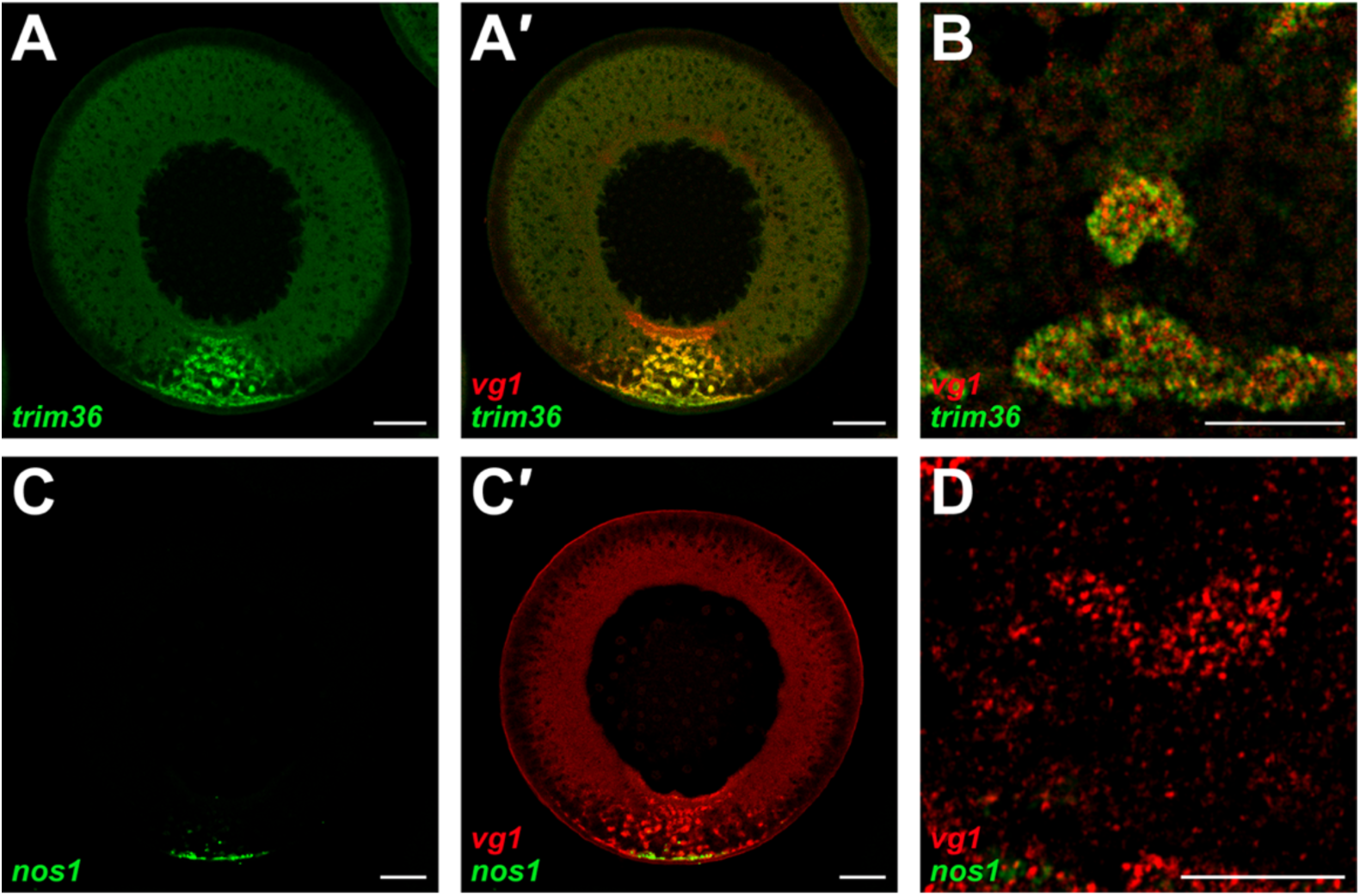
The germinal granule mRNA, nos1, is localized in early oogenesis and is not present in L-bodies. *Trim36* mRNA, which is transported to the vegetal hemisphere during stages II-II of oogenesis is contained in L-bodies, while *nanos1* mRNA, which is found in germ granules and is localized earlier in oogenesis, is not. **(A)** A cryosection of a stage II oocyte probed by FISH for *trim36* mRNA (green), shown merged in A′ with *vg1* mRNA (red) is shown. The vegetal cortex at the bottom. Scale bar=50 μm. **(B)** Higher magnification view of *trim36* mRNA (green) and *vg1* mRNA (red) in the vegetal cytoplasm of a stage II oocyte. Scale bar=10 μm. **(C)** A cryosection of a stage II oocyte probed by FISH for *nanos1* mRNA (green), shown merged in C′ with *vg1* mRNA (red) is shown. The vegetal cortex at the bottom. Scale bar=50 μm. **(D)** Higher magnification view of *nanos1* mRNA (green) and *vg1* mRNA (red) in the vegetal cytoplasm of a stage II oocyte. Scale bar=10 μm.

**Figure S2.**
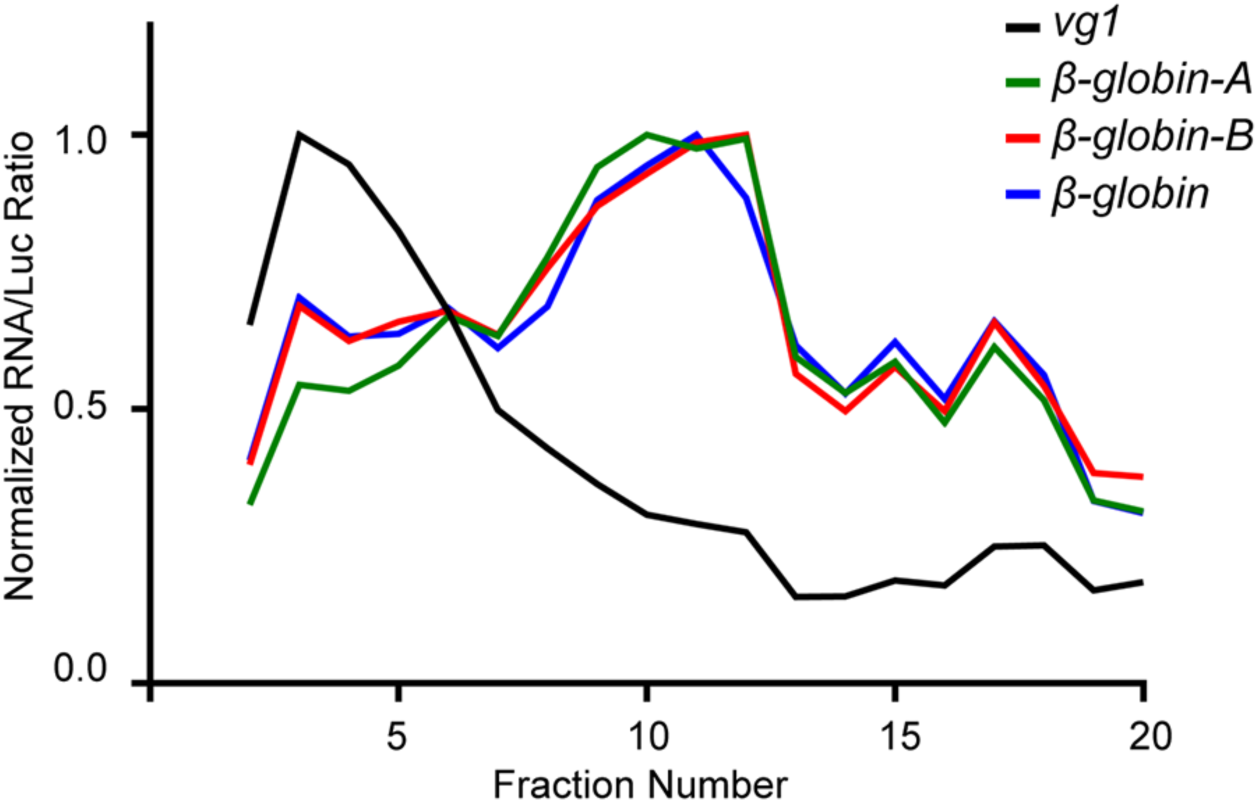
Barcodes used for quantitation do not affect RNP complex assembly or drive incorporation into L-bodies. Because simultaneous detection of the microinjected RNAs by qRT-PCR required tagging of the RNAs with unique barcodes, we tested whether the barcodes used for quantitation of *LE* RNA (LE barcode A) or *mut-VLE* RNA (mutLE barcode B) could affect SE chromatography (as in Fig. 2E) of microinjected *β-globin* RNA complexes. Stage II-III oocytes were injected with *β-globin* RNA (blue, as in Fig. 2E), *β-globin* RNA tagged with LE barcode A (A, green), and *β-globin* RNA tagged with mutLE barcode B (B, red). Oocyte lysates were subjected to SE chromatography and the levels of the microinjected RNAs and endogenous *vg1* mRNA (black) were quantitated in SE column fractions by qRT-PCR, normalized to *luciferase (Luc)* control RNA. While *vg1* RNA (black) was incorporated into large complexes, untagged *β-globin* RNA (blue), LE barcode A (green) tagged *β-globin* RNA, and barcode B (red) tagged *β-globin* RNA were indistinguishable from each other and not incorporated into large RNPs.

**Figure S3.**
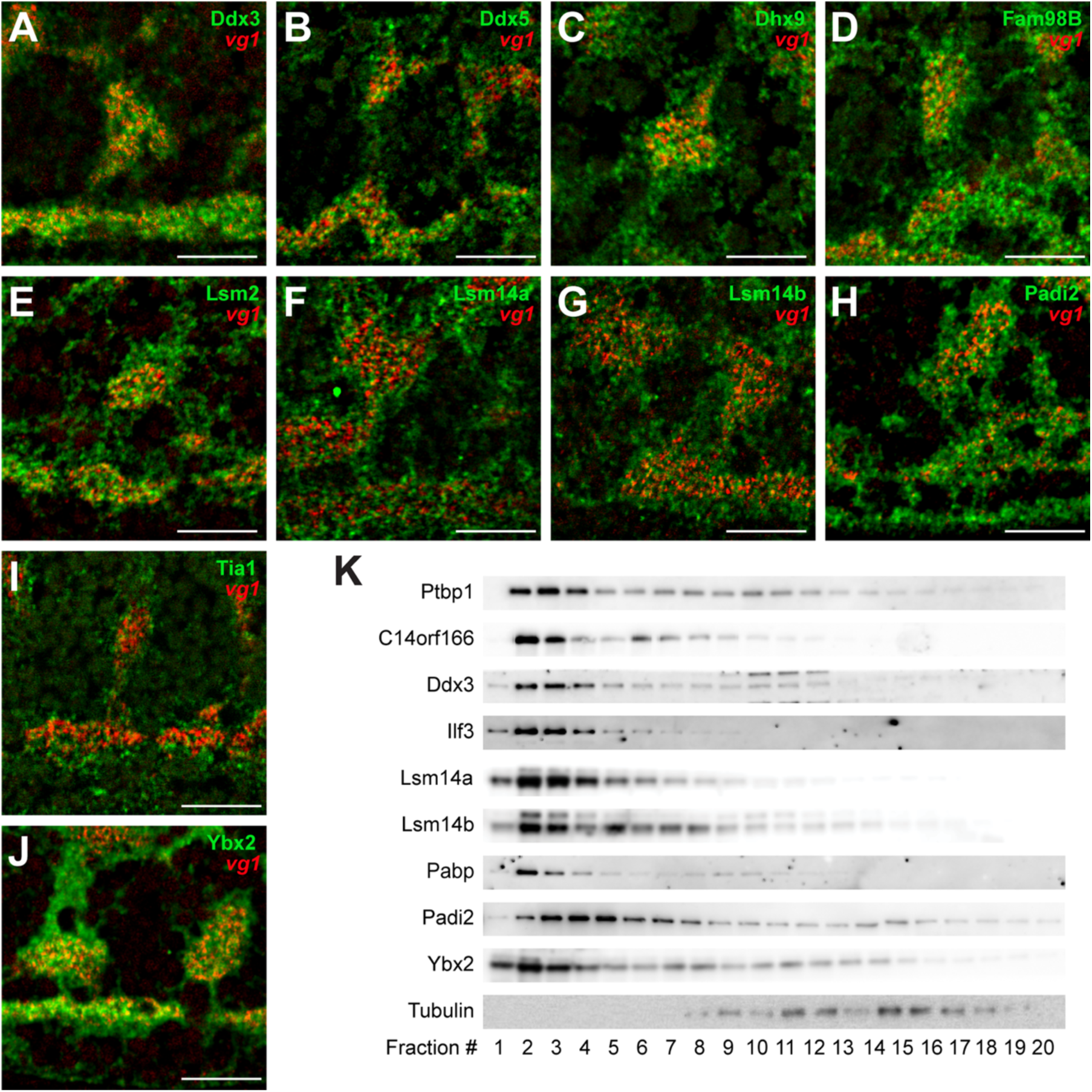
Validation of candidate L-body constituents. Potential L-body constituents were validated by FISH-immunofluorescence (FISH-IF) and size exclusion chromatography. **(A-J)** Shown are FISH-IF images of the vegetal cytoplasm of stage II oocytes with the vegetal cortex at the bottom; scale bars=10μm. Shown in red is *vg1* mRNA detected by FISH. Merged in green is IF using the following antibodies: **(A)** anti-Ddx3, **(B)** anti-Ddx5, **(C)** anti-Dhx9, **(D)** anti-Fam98B, **(E)** anti-Lsm2, **(F)** anti-Lsm14A, **(G)** anti-Lsm14B, **(H)** anti-Padi2, **(I)** anti-Tia1, and **(J)** anti-Ybx2. These potential L-body proteins are all found in large *vg1* mRNA-containing bodies. **(K)** SE column fractions were immunoblotted with the following antibodies: anti-PTB, anti-C14orf166, anti-Ddx3, anti-Ilf3, anti-Lsm14A, anti-Lsm14B, anti-Padi2, anti-Pabp, anti-Ybx2, and anti-Tubulin. The potential L-body proteins, C14orf166, Ddx3, Ilf3, Lsm14A, Lsm14B, Pabp, Padi2 and Ybx2 are enriched in large complexes that chromatograph in the void volume (fractions 1-5), as is the known *vg1* RBP, PTB (Ptbp1). By contrast, Tubulin, is not enriched in the void volume and is present in later fractions. The fraction numbers are shown at the bottom.

**Figure S4.**
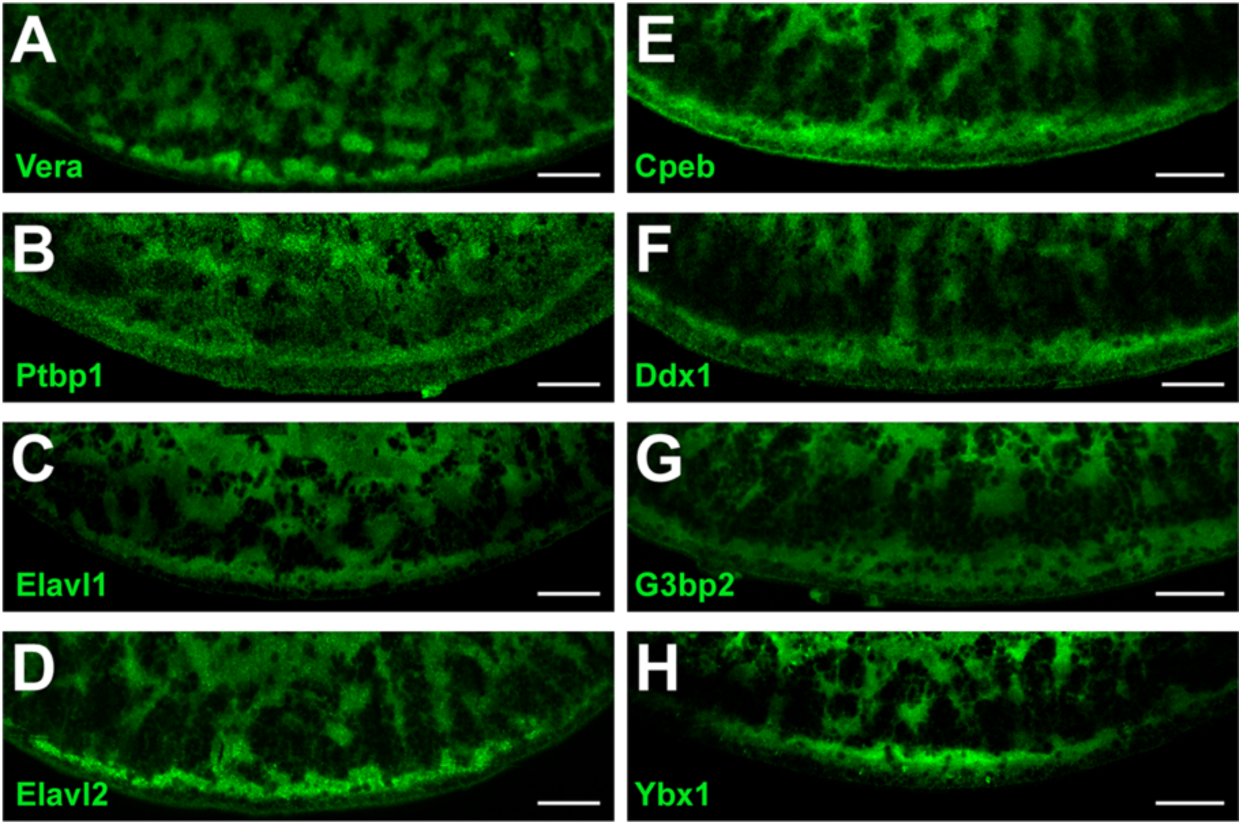
Candidate L-body proteins are enriched in the vegetal cytoplasm. Both known *vg1* RBPs (A-D) and candidate L-body constituents (E-H) are enriched in the vegetal cytoplasm and at the vegetal cortex. Immunofluorescence (IF) was carried out for known *vg1* RBPs using the following antibodies: **(A)** anti-Vera, **(B)** anti-PTB, **(C)** anti-Elav1, and **(D)** anti-Elav2. IF was carried out for potential vegetal RNP transport proteins using the following antibodies: **(E)** anti-Cpeb, **(F)** anti-Ddx1, **(G)** anti-G3bp2, and **(H)** anti-Ybx1. Shown are images of the vegetal cytoplasm of stage II oocytes, with the vegetal cortex at the bottom. Scale bars=20μm.

**Figure S5.**
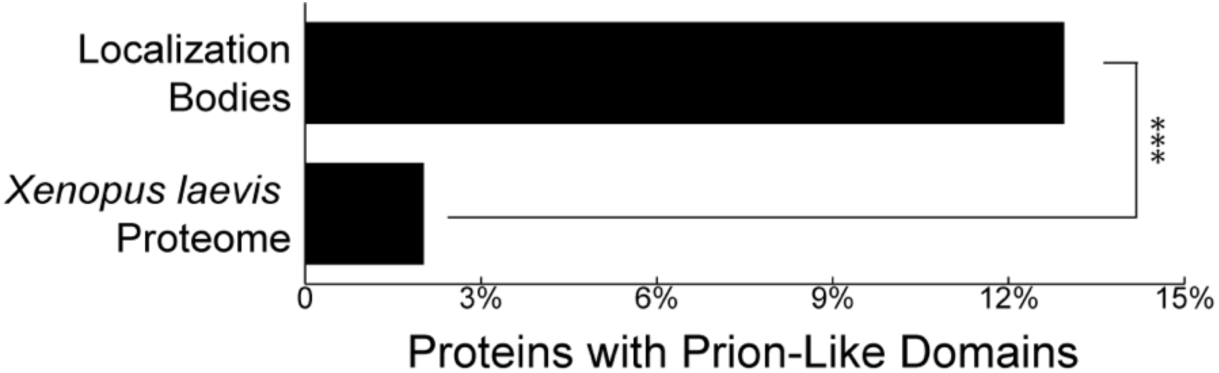
Proteins with prion-like domains are enriched in the L-body proteome. The percentage of prion-like domains in the L-body proteome was compared to the *Xenopus* proteome using PLAAC. Prion-like domains are 6-fold enriched in L-bodies (12.9%) relative to the *Xenopus* proteome (2.02%). *** indicates p < 0.001.

**Figure S6.**
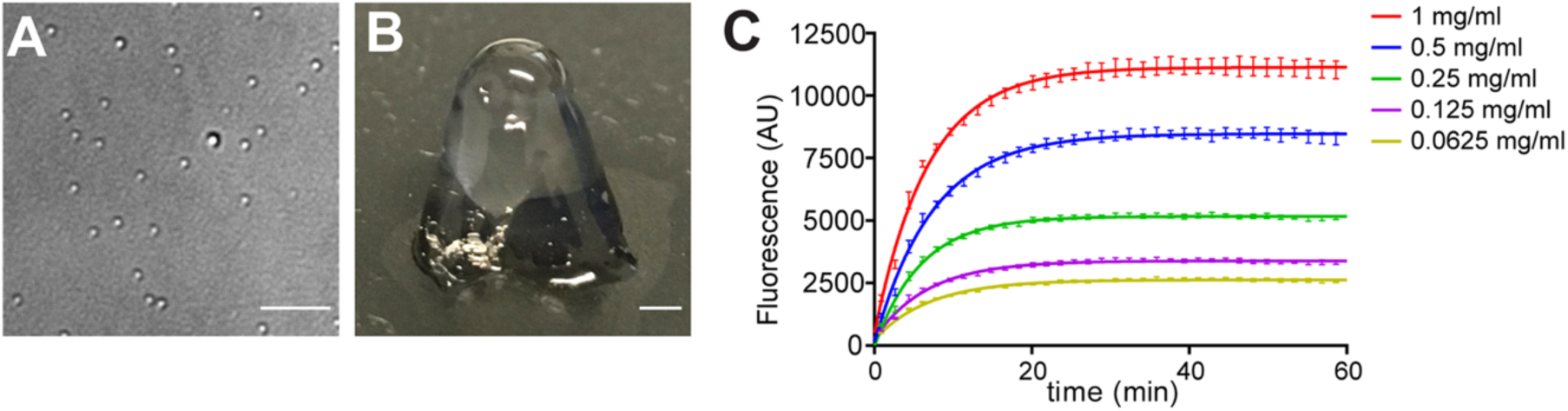
Xenopus oocyte RNA stains with thioflavin and phase-separates in vitro. **(A)** DIC Image of droplets formed at 1 mg/mL oocyte RNA. Scale bar=5μm. **(B)** Image of RNA gel formed at 10 mg/mL *Xenopus* oocyte RNA. Scale bar=1mm. **(C)** Stage II-III *Xenopus* oocyte RNA was serially diluted into buffer containing 125 nM thioflavin-T (in 10 mM Tris pH=7.4, 4 mM MgCl_2_). After heating to denature RNA secondary structure, thioflavin fluorescence was monitored at 482 nm. RNA concentrations were as follows: 0.0625 (yellow), 0.125 (purple), 0.25 (green), 0.5 (blue), and 1 (red) mg/mL.

**Figure S7.**
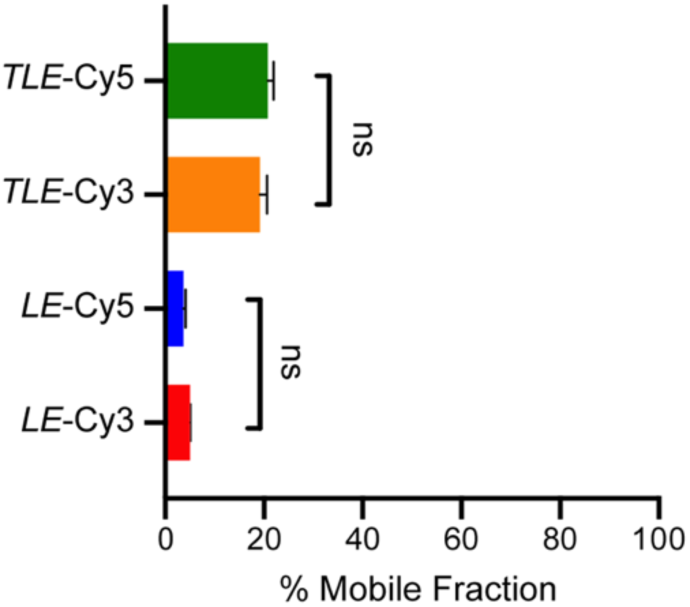
Dynamics of Cy3- and Cy5-labeled RNAs are indistinguishable. Stage II oocytes were microinjected with Cy3-labeled *LE* RNA, Cy5-labeled *LE* RNA, Cy3-labeled *TLE* RNA, or Cy5-labeled *TLE* RNA. FRAP was performed as in Fig. 6. The mobile fractions for Cy3- and Cy5-labled *LE* RNA were 5.05 +/- 0.18 % and 3.67 +/- 0.42 %, respectively. The mobile fractions for Cy3- and Cy5-labled *TLE* RNA were 19.24 +/- 1.41% and 20.82 +/- 1.18%, respectively. The results from 7 oocytes per RNA are shown and error bars indicate standard error of the mean. P-values were calculated using one-way ANOVA with Tukey’s multiple comparison correction; ns is not significant.

## Supplemental Tables

**Table S1.** Validation of candidate L-body constituents. Proteins listed in Fig. 4A, for which antibodies were available, were validated by three approaches: 1) Combined immunofluorescence and FISH for *vg1* RNA (FISH-IF). 2) Chromatography as large complexes using size exclusion chromatography and immunoblot analysis, as in Fig. 2F. 3) Sensitivity of the large complexes to RNase treatment prior to size exclusion chromatography, such that that the large complexes either shift to a smaller size or come out of solution.

| Protein | FISH-IF | Chromatography | RNase Sensitivity |
| --- | --- | --- | --- |
| C14orf166 | + | + | + |
| Cct6 | + | nd | nd |
| Cnot1 | + | nd | nd |
| Cpeb1 | + | + | + |
| Ddx1 | + | + | + |
| Ddx3 | + | + | + |
| Ddx5 | + | + | - |
| Ddx9 | + | nd | nd |
| Dynein | + | nd | nd |
| Eif4g | nd | + | - |
| Elavl1 (HuR) | + | nd | nd |
| Elavl2 (HuB) | + | nd | nd |
| Fam98a | + | nd | nd |
| Fam98b | + | nd | nd |
| Fmrp | + | nd | nd |
| G3bp2 | + | + | + |
| Hnrnpab (40LoVe) | + | + | + |
| Hnrnpd (Auf1) | + | nd | nd |
| Igf2bp3 (Vera) | + | + | + |
| Ilf2 | nd | + | + |
| Ilf3 | + | + | + |
| Lsm1 | na | + | nd |
| Lsm2 | + | nd | nd |
| Lsm14a | + | + | + |
| Lsm14b | + | + | + |
| Nudt16 | + | + | nd |
| Pabp | nd | + | + |
| Padi2 | + | + | nd |
| Prkr | nd | + | + |
| Ptbp1 (PTB) | + | + | + |
| Ruvbl1 | + | nd | nd |
| Stau1 | + | + | + |
| Tia1 | + | nd | nd |
| Ybx1 | + | + | + |
| Ybx2 | + | + | + |

**Table S2.** Gene ontology (GO) analysis of the proteins enriched in L-bodies. The “Count” column indicates the number of proteins identified in Fig. 4A with a given molecular function gene ontology category. The “Percentage” column indicates the percent of proteins in Fig. 4A containing a given gene ontology category. The “P-value” column represents the significance of a measure of overrepresentation of each of these categories, as calculated by DAVID (Huang et al., 2009a; Huang et al., 2009b).

| **Molecular Function – Gene Ontology** | **Count** | **Percentage** | **P-value** |
| --- | --- | --- | --- |
| RNA binding | 27 | 31.4 | 6.46E-19 |
| Nucleotide binding | 18 | 20.9 | 3.96E-11 |
| ATP binding | 15 | 17.4 | 7.78E-02 |
| Nucleic acid binding | 10 | 11.6 | 4.53E-02 |
| mRNA binding | 6 | 7.0 | 5.59E-05 |
| mRNA 3'UTR binding | 4 | 4.7 | 1.59E-03 |
| ATP-dependent RNA helicase activity | 4 | 4.7 | 1.14E-02 |
| Double-stranded RNA binding | 4 | 4.7 | 3.93E-04 |
| Helicase activity | 3 | 3.5 | 2.60E-02 |
| Poly(A) binding | 3 | 3.5 | 1.09E-03 |
| Poly(U) binding | 2 | 2.3 | 2.58E-02 |
| mRNA 5'UTR binding | 2 | 2.3 | 4.23E-02 |

**Table S3.**
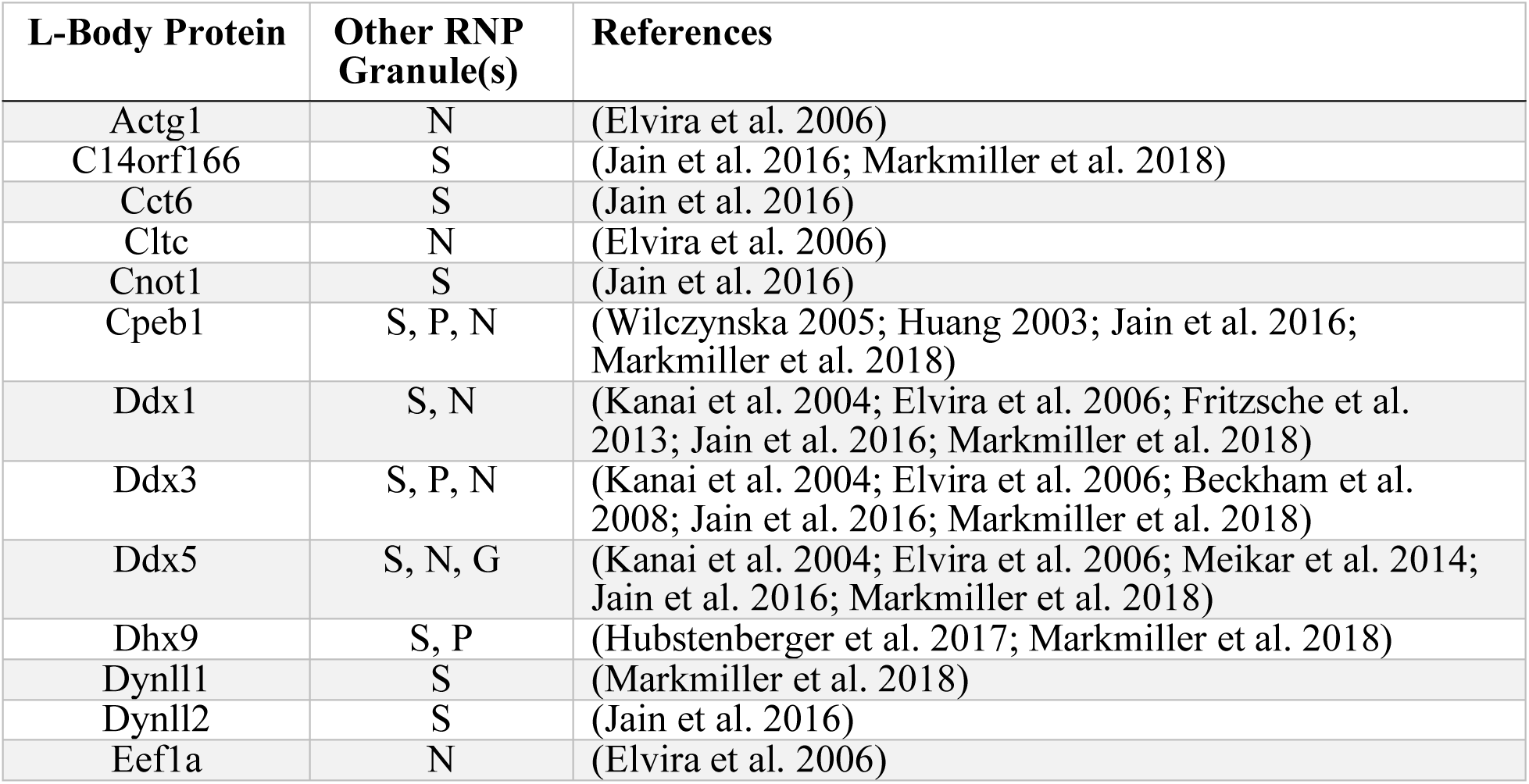
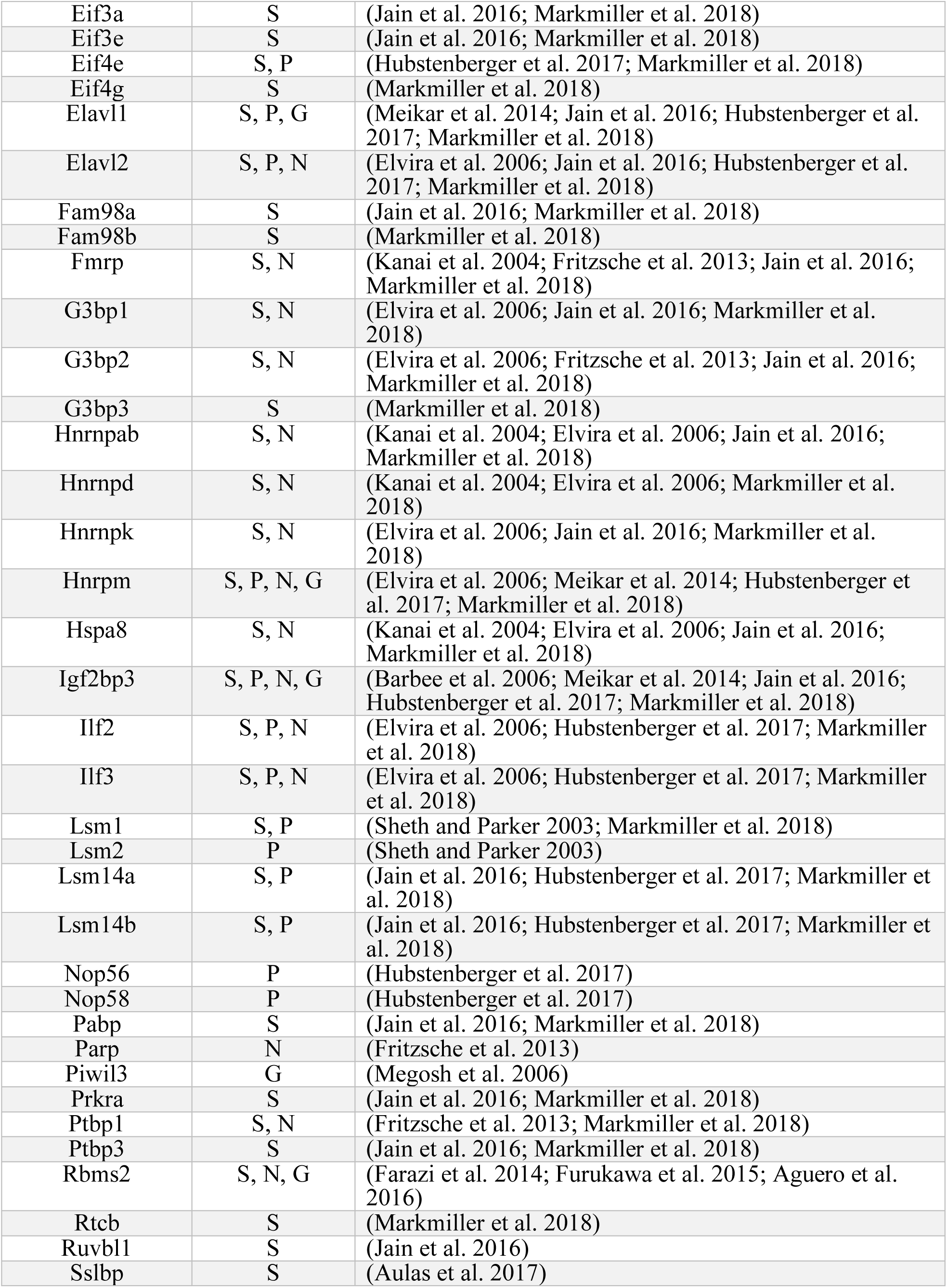
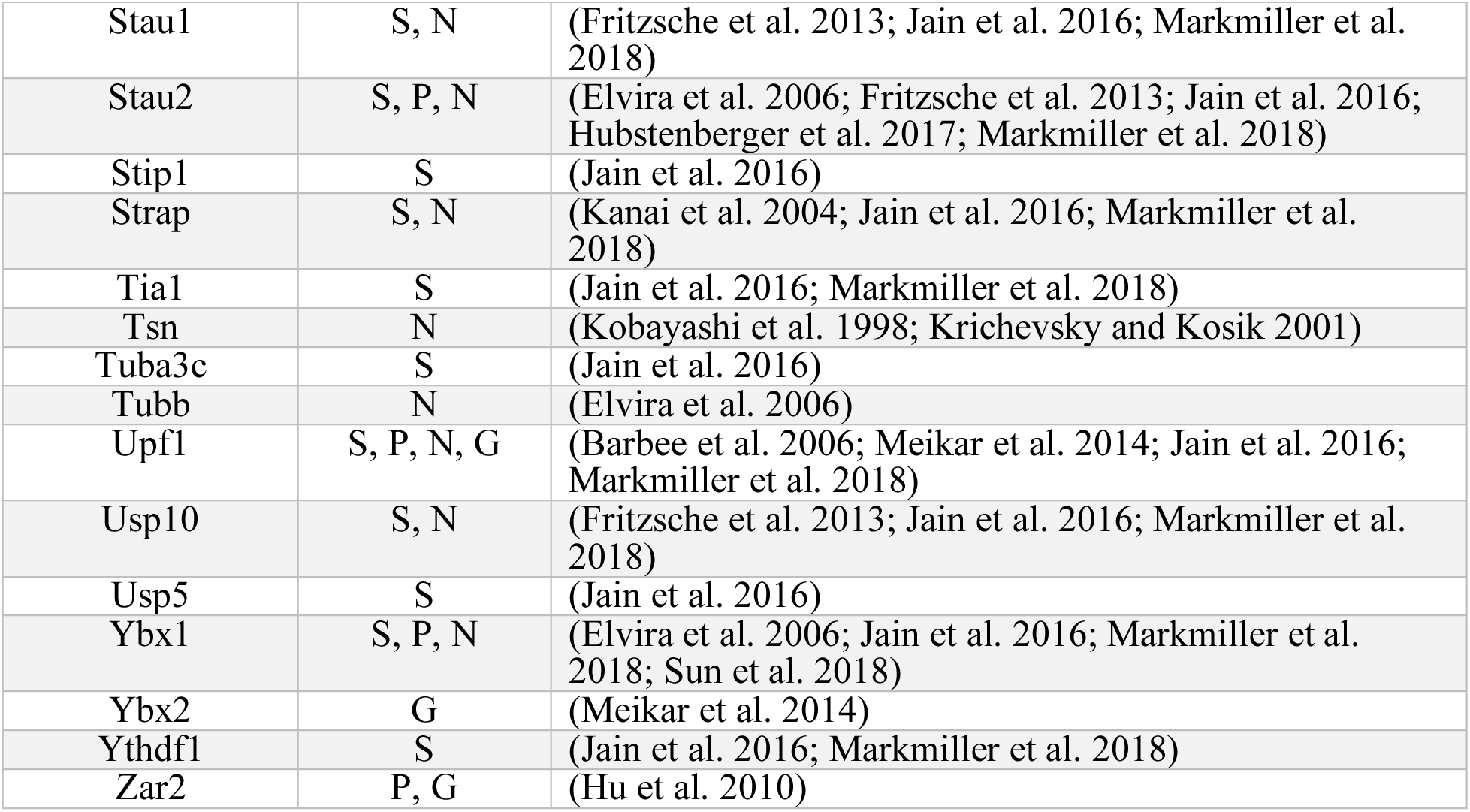
L-body proteins are found in other cytoplasmic RNP granules. L-body proteins (Fig. 4A) that are found in other classes of cytoplasmic RNP granules are listed. The other cytoplasmic granule types are as follows: stress granules (S), P-bodies (P), neuronal granules (N), and germinal granules (G).

**Table S4.**
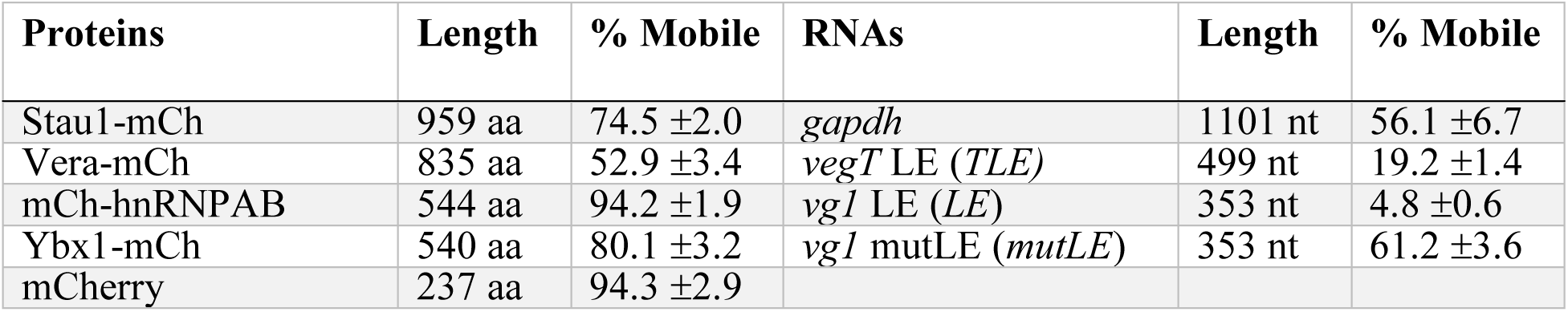
Mobility does not correlate with length of either proteins or RNAs in L-bodies. The lengths of proteins (left) in amino acids (aa) and RNAs (right) in nucleotides (nt) are shown, along with the mobile fractions (% Mobile) determined by the FRAP experiments shown in Fig. 6.

## References

Aguero, T., Zhou, Y., Kloc, M., Chang, P., Houliston, E., and King, M. (2016). Hermes (Rbpms) is a critical component of RNP complexes that sequester germline RNAs during oogenesis. J. Dev. Biol. 4, 2.

Ahsan, N., Belmont, J., Chen, Z., Clifton, J.G., and Salomon, A.R. (2017). Highly reproducible improved label-free quantitative analysis of cellular phosphoproteome by optimization of LC-MS/MS gradient and analytical column construction. J. Proteomics 165, 69–74.

Aulas, A., Fay, M.M., Lyons, S.M., Achorn, C.A., Kedersha, N., Anderson, P., and Ivanov, P. (2017). Stress-specific differences in assembly and composition of stress granules and related foci. J. Cell Sci. 130, 927–937.

Banani, S.F., Rice, A.M., Peeples, W.B., Lin, Y., Jain, S., Parker, R., and Rosen, M.K. (2016). Compositional control of phase-separated cellular bodies. Cell 166, 651–663.

Banani, S.F., Lee, H.O., Hyman, A.A., and Rosen, M.K. (2017). Biomolecular condensates: organizers of cellular biochemistry. Nat. Rev. Mol. Cell Biol. 18, 285–298.

Barbee, S.A., Estes, P.S., Cziko, A.-M., Hillebrand, J., Luedeman, R.A., Coller, J.M., Johnson, N., Howlett, I.C., Geng, C., Ueda, R., Brand, A. H., Newbury, S. F., Wilhelm, J. E., Levine, R. B., Nakamura, A., Parker, R., and Ramaswami, M. (2006). Staufen- and FMRP-containing neuronal RNPs are structurally and functionally related to somatic P bodies. Neuron 52, 997–1009.

Batish, M., van den Bogaard, P., Kramer, F.R., and Tyagi, S. (2012). Neuronal mRNAs travel singly into dendrites. Proc. Natl. Acad. Sci. U. S. A. 109, 4645–4650.

Becker, B.E., and Gard, D.L. (2006). Visualization of the cytoskeleton in Xenopus oocytes and eggs by confocal immunofluorescence microscopy. Methods Mol. Biol. 322, 69–86.

Beckham, C., Hilliker, A., Cziko, A.-M., Noueiry, A., Ramaswami, M., and Parker, R. (2008). The DEAD-Box RNA helicase Ded1p affects and accumulates in Saccharomyces cerevisiae P-Bodies. Mol. Biol. Cell 19, 984–993.

Betley, J.N., Heinrich, B., Vernos, I., Sardet, C., Prodon, F., and Deshler, J.O. (2004). Kinesin II mediates Vg1 mRNA transport in Xenopus oocytes. Curr. Biol. 14, 219–224.

Blower, M.D. (2013). Molecular insights into intracellular RNA localization. Int. Rev. Cell Mol. Biol. 302, 1–39.

Brangwynne, C.P., Tompa, P., and Pappu, R. V. (2015). Polymer physics of intracellular phase transitions. Nat. Phys. 11, 899–904.

Bubunenko M., Kress T.L., Vempati U.D., Mowry K.L., and King M.L. (2002). A consensus RNA signal that directs germ layer determinants to the vegetal cortex of Xenopus oocytes. Dev. Biol. 248, 82–92.

Buchan, J.R. (2014). mRNP granules. Assembly, function, and connections with disease. RNA Biol. 11, 1019–1030.

Buxbaum, A.R., Wu, B., and Singer, R.H. (2014). Single -Actin mRNA detection in neurons reveals a mechanism for regulating its translatability. Science 343, 419–422.

Cote, C. A., Gautreau, D., Denegre, J.M., Kress, T.L., Terry, N. A., and Mowry, K.L. (1999). A Xenopus protein related to hnRNP I has a role in cytoplasmic RNA localization. Mol. Cell 4, 431–437.

Czaplinski, K., Köcher, T., Schelder, M., Segref, A., Wilm, M., and Mattaj, I.W. (2005). Identification of 40LoVe, a Xenopus hnRNP D family protein involved in localizing a TGF-beta-related mRNA during oogenesis. Dev. Cell 8, 505–515.

De Neve J., Thas O., Ottoy J.P., and Clement L. (2013). An extension of the Wilcoxon-Mann-Whitney test for analyzing RT-qPCR data. Stat. Appl. Genet. Mol. Biol. 12, 333–346.

Deshler, J.O., Highett, M.I., and Schnapp, B.J. (1997). Localization of Xenopus Vg1 mRNA by Vera protein and the endoplasmic reticulum. Science 276, 1128–1131.

Elbaum-Garfinkle, S., Kim, Y., Szczepaniak, K., Chen, C.C.-H., Eckmann, C.R., Myong, S., and Brangwynne, C.P. (2015). The disordered P granule protein LAF-1 drives phase separation into droplets with tunable viscosity and dynamics. Proc. Natl. Acad. Sci. U. S. A. 112, 7189–7194.

Elias, J.E., and Gygi, S.P. (2007). Target-decoy search strategy for increased confidence in large-scale protein identifications by mass spectrometry. Nat. Methods 4, 207–214.

Eliscovich, C., Buxbaum, A.R., Katz, Z.B., and Singer, R.H. (2013). mRNA on the move: the road to its biological destiny. J. Biol. Chem. 288, 20361–20368.

Elvira, G., Wasiak, S., Blandford, V., Tong, X.-K., Serrano, A., Fan, X., del Rayo Sánchez-Carbente, M., Servant, F., Bell, A.W., Boismenu, D., Lacaille, J.C., McPherson, P.S., DesGroseillers, L., and Sossin, W.S. (2006). Characterization of an RNA granule from developing brain. Mol. Cell. Proteomics 5, 635–651.

Farazi, T.A., Leonhardt, C.S., Mukherjee, N., Mihailovic, A., Li, S., Max, K.E.A., Meyer, C., Yamaji, M., Cekan, P., Jacobs, N.C., Gerstberger, S., Bognanni, C., Larsson, E., Ohler, U., and Tuschl, T. (2014). Identification of the RNA recognition element of the RBPMS family of RNA-binding proteins and their transcriptome-wide mRNA targets. RNA 20, 1090–1102.

Ferrandon, D. (1997). RNA-RNA interaction is required for the formation of specific bicoid mRNA 3’ UTR-STAUFEN ribonucleoprotein particles. EMBO J. 16, 1751–1758.

Fritzsche, R., Karra, D., Bennett, K.L., Ang, F.Y., Heraud-Farlow, J.E., Tolino, M., Doyle, M., Bauer, K.E., Thomas, S., Planyavsky, M., Arn, E., Bakosova, A., Jungwirth, K., Hörmann, A., Palfi, Z., Sandholzer, J., Schwarz, M., Macchi, P., Colinge, J., Superti-Furga, G., and Kiebler, M.A. (2013). Interactome of two diverse RNA granules links mRNA localization to translational repression in neurons. Cell Rep. 5, 1749–1762.

Furukawa, M.T., Sakamoto, H., and Inoue, K. (2015). Interaction and colocalization of HERMES/RBPMS with NonO, PSF, and G3BP1 in neuronal cytoplasmic RNP granules in mouse retinal line cells. Genes to Cells 20, 257–266.

Gagnon, J.A, and Mowry, K.L. (2011). Molecular motors: directing traffic during RNA localization. Crit. Rev. Biochem. Mol. Biol. 46, 229–239.

Gagnon, J.A., Kreiling, J.A, Powrie, E.A., Wood, T.R., and Mowry, K.L. (2013). Directional transport is mediated by a Dynein-dependent step in an RNA localization pathway. PLoS Biol 11, e1001551.

Gautreau, D., Cote, C.A., and Mowry, K.L. (1997). Two copies of a subelement from the Vg1 RNA localization sequence are sufficient to direct vegetal localization in Xenopus oocytes. Development 124, 5013–5020.

Gomes, E., and Shorter, J. (2019). The molecular language of membraneless organelles. J. Biol. Chem. 294, 7115–7127.

Han, T.W., Kato, M., Xie, S., Wu, L.C., Mirzaei, H., Pei, J., Chen, M., Xie, Y., Allen, J., Xiao, G., and McKnight, S.L. (2012). Cell-free formation of RNA granules: bound RNAs identify features and components of cellular assemblies. Cell 149, 768–779.

Havin, L., Git, A., Elisha, Z., Oberman, F., Yaniv, K., Schwartz, S.P., Standart, N., and Yisraeli, J.K. (1998). RNA-binding protein conserved in both microtubule- and microfilament-based RNA localization. Genes Dev. 12, 1593–1598.

Heasman, J., Wessely, O., Langland, R., Craig, E.J., and Kessler, D.S. (2001). Vegetal localization of maternal mRNAs is disrupted by VegT depletion. Dev. Biol. 240, 377–386.

Holt, C.E., and Schuman, E.M. (2013). The central dogma decentralized: New perspectives on RNA function and local translation in neurons. Neuron 80, 648–657.

Hu, J., Wang, F., Zhu, X., Yuan, Y., Ding, M., and Gao, S. (2010). Mouse ZAR1-Like (XM-359149) colocalizes with mRNA processing components and its dominant-negative mutant caused two-cell-stage embryonic arrest. Dev. Dyn. 239, 407–424.

Huang, D.W., Sherman, B.T., and Lempicki, R.A. (2009a). Systematic and integrative analysis of large gene lists using DAVID bioinformatics resources. Nat. Protoc. 4, 44–57.

Huang, D.W., Sherman, B.T., and Lempicki, R.A. (2009b). Bioinformatics enrichment tools: paths toward the comprehensive functional analysis of large gene lists. Nucleic Acids Res. 37, 1–13.

Huang, Y.S., Carson, J.H., Barbarese, E., and Richter, J.D. (2003). Facilitation of dendritic mRNA transport by CPEB. Genes Dev. 17, 638–653.

Hubstenberger, A., Noble, S.L., Cameron, C., and Evans, T.C. (2013). Translation repressors, an RNA helicase, and developmental cues control RNP phase transitions during early development. Dev. Cell 27, 161–173.

Hubstenberger, A., Cameron, C., Noble, S.L., Keenan, S., and Evans, T.C. (2015). Modifiers of solid RNP granules control normal RNP dynamics and mRNA activity in early development. J. Cell Biol. 211, 703–716.

Hubstenberger, A., Courel, M., Bénard, M., Souquere, S., Ernoult-Lange, M., Chouaib, R., Yi, Z., Morlot, J.-B., Munier, A., Fradet, M., et al. (2017). P-Body purification reveals the condensation of repressed mRNA regulons. Mol. Cell 68, 144–157.

Jain, A., and Vale, R.D. (2017). RNA phase transitions in repeat expansion disorders. Nature 546, 243–247.

Jain, S., Wheeler, J.R., Walters, R.W., Agrawal, A., Barsic, A., and Parker, R. (2016). ATPase-modulated stress granules contain a diverse proteome and substructure. Cell 164, 487–498.

Jambor, H., Brunel, C., and Ephrussi, A. (2011). Dimerization of oskar 3’ UTRs promotes hitchhiking for RNA localization in the Drosophila oocyte. RNA 17, 2049–2057.

Kanai, Y., Dohmae, N., and Hirokawa, N. (2004). Kinesin transports RNA: isolation and characterization of an RNA-transporting granule. Neuron 43, 513–525.

Kato, M., Han, T.W., Xie, S., Shi, K., Du, X., Wu, L.C., Mirzaei, H., Goldsmith, E.J., Longgood, J., Pei, J., Grishin, N.V., Frantz, D.E., Schneider, J.W., Chen, S., Li, L., Sawaya, M.R., Eisenberg, D., Tycko, R., and McKnight, S.L. (2012). Cell-free formation of RNA granules: low complexity sequence domains form dynamic fibers within hydrogels. Cell 149, 753–767.

Kedersha, N., Ivanov, P., and Anderson, P. (2013). Stress granules and cell signaling: more than just a passing phase? Trends Biochem. Sci. 38, 494–506.

Kobayashi, S., Takashima, A., and Anzai, K. (1998). The dendritic translocation of translin protein in the form of BC1 RNA protein particles in developing rat hippocampal neurons in primary culture. Biochem. Biophys. Res. Commun. 253, 448–453.

Krichevsky, A.M., and Kosik, K.S. (2001). Neuronal RNA granules: A link between RNA localization and stimulation-dependent translation. Neuron 32, 683–696.

Krieg, P.A., and Melton, D.A. (1984). Functional messenger RNAs are produced by SP6 in vitro transcription of cloned cDNAs. Nucleic Acids Res. 12, 7057–7070.

Lancaster, A.K., Nutter-Upham, A., Lindquist, S., and King, O.D. (2014). PLAAC: a web and command-line application to identify proteins with prion-like amino acid composition. Bioinformatics 30, 2501–2502.

Langdon, E.M., Qiu, Y., Ghanbari Niaki, A., McLaughlin, G.A., Weidmann, C.A., Gerbich, T.M., Smith, J.A., Crutchley, J.M., Termini, C.M., Weeks, K.M., Myong, S., and Gladfelter, A.S. (2018). mRNA structure determines specificity of a polyQ-driven phase separation. Science 360, 922–927.

Lewis, R.A., Kress, T.L., Cote, C.A., Gautreau, D., Rokop, M.E., and Mowry, K.L. (2004). Conserved and clustered RNA recognition sequences are a critical feature of signals directing RNA localization in Xenopus oocytes. Mech Dev. 121, 101–109.

Lewis, R.A., Gagnon, J.A., and Mowry, K.L. (2008). PTB/hnRNP I is required for RNP remodeling during RNA localization in Xenopus oocytes. Mol. Cell. Biol. 28, 678–686.

Li, P., Banjade, S., Cheng, H.-C., Kim, S., Chen, B., Guo, L., Llaguno, M., Hollingsworth, J. V., King, D.S., Banani, S.F., Russo, P.S., Jiang, Q., Nixon, B.T., and Rosen, M.K. (2012). Phase transitions in the assembly of multivalent signalling proteins. Nature 483, 336–340.

Lin, Y., Protter, D.S.W., Rosen, M.K., and Parker, R. (2015). Formation and maturation of phase-separated liquid droplets by RNA-binding proteins. Mol. Cell 60, 208–219.

Little, S.C., Sinsimer, K.S., Lee, J.J., Wieschaus, E.F., and Gavis, E.R. (2015). Independent and coordinate trafficking of single Drosophila germ plasm mRNAs. Nat. Cell Biol. 17, 558–568.

Markmiller, S., Soltanieh, S., Server, K.L., Mak, R., Jin, W., Fang, M.Y., Luo, E.C., Krach, F., Yang, D., Sen, A., Fulzele, A., Wozniak, J.M., Gonzalez, D.J., Kankel, M.W., Gao, F.B., Bennett, E.J., Lécuyer, E., and Yeo, G.W. (2018). Context-dependent and disease-specific diversity in protein interactions within stress granules. Cell 172, 590–604.

Medioni, C., Mowry, K., and Besse, F. (2012). Principles and roles of mRNA localization in animal development. Development 139, 3263–3276.

Megosh, H.B., Cox, D.N., Campbell, C., and Lin, H. (2006). The role of PIWI and the miRNA machinery in Drosophila germline determination. Curr. Biol. 16, 1884–1894.

Meikar, O., Vagin, V. V., Chalmel, F., Sõstar, K., Lardenois, A., Hammell, M., Jin, Y., Da Ros, M., Wasik, K.A., Toppari, J., Hannon, G.J., and Kotaja, N. (2014). An atlas of chromatoid body components. RNA 20, 483–495.

Messitt, T.J., Gagnon, J.A., Kreiling, J.A., Pratt, C.A., Yoon, Y.J., and Mowry, K.L. (2008). Multiple kinesin motors coordinate cytoplasmic RNA transport on a subpopulation of microtubules in Xenopus oocytes. Dev. Cell 15, 426–436.

Miura, K., Rueden, C., Hiner, M., Schindelin, J., and Rietdorf, J. (2014). ImageJ Plugin CorrectBleach V2.0.2 (Zenodo). http://dx.doi.org/10.5281/zenodo.596358.

Neil, C.R., and Mowry, K. (2018). Fluorescence in situ hybridization of cryosectioned Xenopus oocytes. Cold Spring Harb. Protoc. pdb.prot097030.

Niepielko, M.G., Eagle, W.V.I., and Gavis, E.R. (2018). Stochastic seeding coupled with mRNA self-recruitment generates heterogeneous Drosophila germ granules. Curr. Biol. 28, 1872–1881.e3.

Nilsson, M.R. (2004). Techniques to study amyloid fibril formation in vitro. Methods 34, 151–160.

Nott, T.J., Petsalaki, E., Farber, P., Jervis, D., Fussner, E., Plochowietz, A., Craggs, T.D., Bazett-Jones, D.P., Pawson, T., Forman-Kay, J.D., and Baldwin, A.J. (2015). Phase transition of a disordered nuage protein generates environmentally responsive membraneless organelles. Mol. Cell 57, 936–947.

Peng, Z., Mizianty, M.J., and Kurgan, L. (2014). Genome-scale prediction of proteins with long intrinsically disordered regions. Proteins Struct. Funct. Bioinforma. 82, 145–158.

Perez-Riverol, Y., Csordas, A., Bai, J., Bernal-Llinares, M., Hewapathirana, S., Kundu, D.J., Inuganti, A., Griss, J., Mayer, G., Eisenacher, M., Pérez, E., Uszkoreit, J., Pfeuffer, J., Sachsenberg, T., Yilmaz, Ş., Tiwary, S., Cox, J., Audain, E., Walzer, M., Jarnuczak, A.F., Ternent, T., Brazma, A., and Vizcaíno, J.A. (2019). The PRIDE database and related tools and resources in 2019: Improving support for quantification data. Nucleic Acids Res. 47, 442–450.

Pfaffl, M.W. (2001). A new mathematical model for relative quantification in real-time RT-PCR. Nucleic Acids Res. 29, e45.

Protter, D.S.W., Rao, B.S., Van Treeck, B., Lin, Y., Mizoue, L., Rosen, M.K., and Parker, R. (2018). Intrinsically disordered regions can contribute promiscuous interactions to RNP granule assembly. Cell Rep. 22, 1401–1412.

Putnam, A., Cassani, M., Smith, J., and Seydoux, G. (2019). A gel phase promotes condensation of liquid P granules in Caenorhabditis elegans embryos. Nat. Struct. Mol. Biol. 26, 220–226.

Ramakers, C., Ruijter, J.M., Deprez, R.H.L., and Moorman, A.F.M. (2003). Assumption-free analysis of quantitative real-time polymerase chain reaction (PCR) data. Neurosci. Lett. 339, 62–66.

Rebagliati, M.R., Weeks, D.L., Harvey, R.P., and Melton, D.A. (1985). Identification and cloning of localized maternal RNAs from Xenopus eggs. Cell 42, 769–777.

Schisa, J.A. (2012). New insights into the regulation of RNP granule assembly in oocytes. Int. Rev. Cell Mol. Biol. 295, 233–289.

Sheth, U., and Parker R. (2003). Decapping and decay of messenger RNA occur in cytoplasmic processing bodies. Science 300, 805–808.

Shin, Y., and Brangwynne, C.P. (2017). Liquid phase condensation in cell physiology and disease. Science 357, eaaf4382.

Sun, J., Yan, L., Shen, W., and Meng, A. (2018). Maternal Ybx1 safeguards zebrafish oocyte maturation and maternal-to-zygotic transition by repressing global translation. Development 145, dev166587.

Suter, B. (2018). RNA localization and transport. Biochim. Biophys. Acta - Gene Regul. Mech. 1861, 938–951.

Tafuri, S.R., and Wolffe, A.P. (1990). Xenopus Y-box transcription factors: Molecular cloning, functional analysis, and developmental regulation. Proc. Natl. Acad. Sci. U. S. A. 87, 9028–9032.

Toretsky, J.A., and Wright, P.E. (2014). Assemblages: functional units formed by cellular phase separation. J. Cell Biol. 206, 579–588.

Van Treeck, B., Protter, D.S.W., Matheny, T., Khong, A., Link, C.D., and Parker, R. (2018). RNA self-assembly contributes to stress granule formation and defining the stress granule transcriptome. Proc. Natl. Acad. Sci. U. S. A. 115, 2734–2739.

Wang, J.T., Smith, J., Chen, B.-C., Schmidt, H., Rasoloson, D., Paix, A., Lambrus, B.G., Calidas, D., Betzig, E., and Seydoux, G. (2014). Regulation of RNA granule dynamics by phosphorylation of serine-rich, intrinsically disordered proteins in C. elegans. Elife 3, e04591.

Weber, S.C., and Brangwynne, C.P. (2012). Getting RNA and protein in phase. Cell 149, 1188–1191.

Weeks, D.L., and Melton, D.A. (1987). A maternal mRNA localized to the vegetal hemisphere in Xenopus eggs codes for a growth factor related to TGF-beta. Cell 51, 861–867.

Wilczynska, A., Aigueperse, C., Kress, M., Dautry, F., and Weil, D. (2005). The translational regulator CPEB1 provides a link between dcp1 bodies and stress granules. J. Cell Sci. 118, 981–992.

Xu, S., Li, Q., Xiang, J., Yang, Q., Sun, H., Guan, A., Wang, L., Liu, Y., Yu, L., Shi, Y., Chen, H., and Tang, Y. (2016). Thioflavin T as an efficient fluorescence sensor for selective recognition of RNA G-quadruplexes. Sci. Rep. 6, 24793.

Yamazaki, T., Souquere, S., Chujo, T., Kobelke, S., Chong, Y.S., Fox, A.H., Bond, C.S., Nakagawa, S., Pierron, G., and Hirose, T. (2018). Functional domains of NEAT1 architectural lncRNA induce paraspeckle assembly through phase separation. Mol. Cell 70, 1038–1053.e7.

Yoon, Y.J., and Mowry, K.L. (2004). Xenopus Staufen is a component of a ribonucleoprotein complex containing Vg1 RNA and kinesin. Development 131, 3035–3045.

Yu, K., and Salomon, A.R. (2009). PeptideDepot: Flexible relational database for visual analysis of quantitative proteomic data and integration of existing protein information. Proteomics 9, 5350–5358.

Yu, K., and Salomon, A.R. (2010). HTAPP: High-throughput autonomous proteomic pipeline. Proteomics 10, 2113–2122.

Yu, K., Sabelli, A., DeKeukelaere, L., Park, R., Sindi, S., Gatsonis, C.A., and Salomon, A. (2009). Integrated platform for manual and high-throughput statistical validation of tandem mass spectra. Proteomics 9, 3115–3125.

